# Imaging angiogenesis in an intracerebrally induced model of brain macrometastasis using α_v_β_3_-targeted iron oxide microparticles

**DOI:** 10.1101/2023.03.14.532679

**Authors:** Jessica Buck, Francisco Perez-Balderas, Niloufar Zarghami, Vanessa Johanssen, Alexandre A Khrapitchev, James R Larkin, Nicola R Sibson

## Abstract

Brain metastasis is responsible for a large proportion of cancer mortality, and there are currently no effective treatments. Moreover, the impact of treatments, particularly anti-angiogenic therapeutics, is difficult to ascertain using current magnetic resonance imaging (MRI) methods. Imaging of the angiogenic vasculature has been successfully carried out in solid tumours using microparticles of iron oxide (MPIO) conjugated to a peptide (RGD) targeting the integrin α_v_β_3_. The aim of this study was to determine whether RGD-MPIO could be used to identify angiogenic blood vessels in brain metastases *in vivo.* A mouse model of intracerebrally implanted brain macrometastasis was established through intracerebral injection of 4T1-GFP cells. T_2_* weighted imaging was used to visualize MPIO induced hypointense voxels *in vivo,* and Prussian blue staining was used to visualize MPIO and endogenous iron histologically *ex vivo.* The RGD-MPIO showed target-specific binding *in vivo,* but the sensitivity of the agent for visualizing angiogenic vessels *per se* was reduced by the presence of endogenous iron-laden macrophages in larger metastases, resulting in pre-existing hypointense areas within the tumour. Further, our data suggest that peptide-targeted MPIO, but not antibody-targeted MPIO, are taken up by perivascular macrophages within the macrometastatic microenvironment, resulting in additional nonspecific contrast. Whilst pre-MPIO imaging will circumvent the issues surrounding pre-existing hypointensities and enable detection of specific contrast, our findings suggest that the use of antibodies rather than peptides as the targeting ligand may represent a preferable route forward for new angiogenesis-targeted molecular MRI agents.

## Introduction

Metastasis, or secondary spread from a primary tumour to distant sites, is one of the enduring challenges in the treatment of cancer. The detection and treatment of brain metastases remains an important clinical challenge. Thus, there is a need for the development of targeted therapies to treat brain metastases, together with more accurate strategies for measuring treatment response. Anti-angiogenic agents, predominantly targeting VEGF and VEGFR2, have been and are currently being evaluated in clinical trials for the treatment of human brain metastasis^1^. Angiogenesis is crucial for tumours to grow beyond a few millimetres in size^2^ and, thus, represents an important therapeutic target. Despite showing great promise in initial pre-clinical studies, the use of anti-angiogenic drugs in the clinic has had mixed results. There are many reasons for these discrepancies, including an inability to stratify patients by angiogenesis levels, an inability to monitor treatment response using conventional imaging, and the activation of alternative signalling pathways mediating drug resistance.

Angiogenesis is mediated by many different factors and pathways, one of which is the integrin α_v_β_3_. Also known as the vitronectin receptor, integrin α_v_β_3_ is expressed at high levels on angiogenic endothelial cells^3,4^ and some types of tumour cells^5–7^, but only at very low levels on the normal vasculature. Integrin α_v_β_3_ has a specific binding site in the peptide motif Arg-Gly-Asp (RGD)^8^ and, consequently, RGD peptides are used in a wide range of blocking and targeting applications, including small molecule drugs and molecular imaging. Anti-angiogenic drugs targeting α_v_β_3_ such as Cilengitide and Etaracizumab have shown promise in preclinical studies, but have not been successful in clinical trials, particularly in the brain and in metastatic cancer^9–15^. It is important to note that none of these clinical trials have attempted to stratify patients by α_v_β_3_ expression levels, which might be expected to predict response. In the only trial that attempted to image α_v_β_3_ using radiolabelled Etaracizumab, the imaging was unsuccessful, with only subtle localisation to the tumour bed, which the authors proposed was due to instability of ^99m^Tc labelling^12^. These findings highlight the need for sensitive, specific, and reproducible clinical imaging of the angiogenic vasculature in tumours, and brain metastases in particular.

Molecular MRI strategies that have previously been used to image angiogenesis include the conjugation of anti-α_v_β_3_ antibodies or various RGD peptides to paramagnetic gadolinium containing liposomes^16–20^, perfluorocarbon based nanoparticles^21–24^, or lipoprotein cores^25^. However, the clearance rates, specificity and signal-to-noise ratio (SNR) of these particles remained low, leading to the investigation of targeted iron oxide-based agents. Targeting of ultra-small particles of iron oxide (USPIO) to α_v_β_3_ has been reported, and in some studies with sufficient SNR to be detected on a clinical scanner^26,27^. In preclinical studies, this approach has been suggested to enable differentiation between integrin α_v_β_3_ expressing angiogenic blood vessels and integrin α_v_β_3_ overexpressing tumour cells^26^, as well as prediction or measurement of response to treatment^28,29^. Despite the increased sensitivity compared to gadolinium-based molecular imaging agents, however, the propensity of these nanoparticles to extravasate into tissue remains a significant confound, as the contrast produced is no longer specific to their endothelial target^30–33^. Moreover, nano-sized particles have a relatively long half-life in the circulation (~16.5 hours), which is a limitation to imaging of target-specific binding^33^. In contrast, microparticles of iron oxide (MPIO), are significantly larger than USPIO, exhibit a much shorter blood half-life ~1 min^33–35^, and are obligate intravascular agents. Consequently, MPIO have major advantages over USPIO for molecular imaging of endovascular targets, with greater contrast effects owing to their higher iron content, and high sensitivity for target-specific binding owing to an absence of background contrast effects from circulating particles. For this reason, we have previously developed a cyclic-RGD conjugated MPIO construct (RGD-MPIO) and have demonstrated its potential for imaging angiogenic tumour vasculature in solid tumours^35^.

Thus, in order to address the unmet need for a specific and sensitive method for monitoring tumour angiogenesis, the primary aim of the current study was to determine whether RGD-MPIO in conjunction with MRI enables detection of angiogenic vasculature in an intracerebrally implanted model of brain macrometastases.

## Materials and methods

### Preparation of peptide and antibody conjugated MPIO

Cyclic RGD conjugated MPIO were prepared using 1 μm diameter Dynabead MyOne carboxylic acid MPIO (65011, Fisher Scientific, UK). MPIO were washed in MES buffer and resuspended, then incubated with EDC in a shaking platform. MPIO were washed with MES buffer, then incubated for 24 hours with 10 mM cyclic RGD or control scrambled RDG peptides (Supplementary Figure 1). The MPIO were then washed and stored in PBS at 4°C on a shaking platform.

Antibody conjugated MPIO were prepared as described by Zarghami et al. using 40 μg VCAM-1 (1510-14, Southern Biotech, Birmingham, USA) or IgG (0116-14, Southern Biotech, Birmingham, USA) antibody for 1 mg of 1 μm diameter ProMag carboxylic acid MPIO beads (PMC1N, Stratech, Ely, UK).

### In vitro binding of cyclic RGD peptides to mouse integrin α_v_β_3_

The binding efficiency of various RGD peptides has been systematically measured previously for human integrin α_v_β_3_^36^, but not for mouse integrin α_v_β_3_. Thus, we tested the binding of three different RGD cyclic peptides, and one control scrambled RDG peptide (structures shown in Supplementary Figure 1), to mouse integrin α_v_β_3_. These peptides were conjugated separately to MPIO, as described below.

To measure the binding of RGD-MPIO to mouse integrin α_v_β_3_ under flow conditions, glass capillaries were mounted onto a narrow-gauge syringe with dry-cure glue. First, each glass capillary was filled with a mixture of o-Xylene, Triethoxy-(3-glycidyloxypropyl)silane (TEGOPS) and N,N-Diisopropylethylamine (DIPEA; ratio 1000:320:10), incubated for 17 hours at 80° with phosphorus pentoxide, then washed with o-xylene and methanol, and dried for 4 hours at room temperature under vacuum conditions. Subsequently, capillaries were filled with 200 ng/mL mouse integrin α_v_β_3_ (7889-AV-050, R&D Systems, Abingdon, UK) and incubated at room temperature for 24 hours. Remaining functional groups were quenched with 10 mM ethanolamine for 24 hours at room temperature. Control capillaries were filled with 200 ng/mL BSA and incubated as above. Capillaries were then washed and stored filled with PBS at 4°C.

Functionalised capillaries were mounted under a Nikon TE 2000-U microscope, and RGD-MPIO or control RDG-MPIO were resuspended in 30% sucrose in PBS to mimic the viscosity of blood. The sucrose-MPIO solution was infused through the capillaries, using a syringe pump, at 20 μL/min to mimic hydraulic shear conditions in blood vessels. The volume of solution and infusion time were recorded in order to calculate flow rates retrospectively. Photomicrographs were taken across 8 separate fields-of-view (FOV) along the capillary, and the number of bound MPIO per FOV manually counted.

Blocking experiments were also carried out to test the specificity of binding of RGD-MPIO. HUVEC-C cells were cultured in Media 199 supplemented with 10% fetal calf serum, penicillin and streptomycin (100 μg/mL). Once confluent, cells were stimulated to express integrin α_v_β_3_ by stimulation for 20 hours with S-nitroso-n-acetylpenicillamine (SNAP). Blocking was carried out by adding RGD or control RDG peptide to the media to a final concentration of 10 μM, and incubating for 1 hour. Media was removed, cells rinsed with PBS, and fresh media added. 1 μg of RGD-MPIO or control RDG-MPIO was then added to each well, incubated in a shaking incubator for 40 minutes at 37° and 2500 rpm. Cells were rinsed 5 times with PBS to remove any unbound MPIO, then fixed with 4% paraformaldehyde. Images of 10 FOV per condition were taken using an Olympus IX-71 inverted microscope at 40x magnification, and bound MPIO counted by an observer blinded to condition.

### Animal models

All mice were housed in individually ventilated cages with food and water provided *ad libitum.* All animal experiments were approved by the University of Oxford Clinical Medicine Ethics Review Committee and the UK Home Office (Animals [Scientific Procedures] Act 1986), and conducted in accordance with the University of Oxford Policy on the Use of Animals in Scientific Research, the ARRIVE Guidelines and Guidelines for the Welfare and Use of Animals in Cancer Research^37^.

To study the angiogenic vasculature of brain metastases, an intracerebrally implanted mouse model of breast cancer brain macrometastasis was used. 4T1 tumours naturally metastasise to the brain when a primary tumour is implanted into the mammary fat pad^38,39^, and brain metastases can also be induced haematogenously by intracardiac or intracarotid injection of 4T1 cells, which is considered to be a close representation of the natural route of seeding to the brain. However, intracardiac injection also seeds 4T1 cells to lung, liver and bone, and mice become moribund before macroscopic metastases develop within the brain. Consequently, in order to study larger, neo-vascularised brain metastases, the intracerebral route of induction is necessary. We have previously demonstrated that intracerebral injection of metastatic breast cancer cells leads to tumour growth that is very similar to that found when metastases are induced via intracardiac injection, as well as to human brain metastasis growth^40^.

Female BALB/c mice (6-10 weeks old) were anaesthetized with 3% isoflurane in 30% O_2_: 70% N_2_O and focally microinjected with 5000 4T1-GFP metastatic murine mammary carcinoma cells^41^ into the left striatum. Mice were randomly allocated to timepoint and treatment groups. Following imaging, mice underwent transcardial perfusion-fixation wat weekly timepoints 1-5 weeks after tumour induction; and the brains were collected for histology. Brains were cryoprotected in 30% w/v sucrose, frozen in isopentane and dry ice, and 10 μm cryosections were cut through the striatum.

To induce focal BBB breakdown in one cerebral hemisphere, female BALB/c mice were anaesthetized and injected as described above with 1 μg CINC-1 protein (453-KC-050/CF, R&D systems, Abingdon, UK) in 1 μL saline in the left striatum, as described previously^42^. Mice underwent imaging 5 hours after injection. Following imaging, mice were transcardially perfusion-fixed, and brains collected and sectioned as above.

### Magnetic resonance imaging

4T1-GFP brain macrometastases were induced as described above, and each mouse was imaged once using RGD-MPIO at day 7 (n = 3), 14 (n = 2), 21 (n = 3), 28 (n = 4) or 35 (n = 5). Further mice were also imaged using a control RDG-MPIO at day 7 (n = 3), 14 (n = 3), 21 (n = 3), 28 (n = 4) or 35 (n = 3); and at day 28 using VCAM-MPIO (n = 3) or IgG-MPIO (n = 2). In the CINC-1 model of BBB permeabilisation, mice underwent imaging 5h after CINC-1 injection using RGD-MPIO (n = 3), control RDG-MPIO (n = 3), VCAM-MPIO (n = 3), or IgG-MPIO (n = 3).

Experiments were performed using a 9.4 T Agilent DDR MRI spectrometer (Agilent Technologies, Santa Clara, CA, USA) with a 160 mm horizontal bore, and an internal diameter 100 mm / outer diameter 155 mm shim and gradient system (Agilent Technologies, Santa Clara, CA, USA) with maximum strength of 400 mT/m. A 26 mm internal diameter volume transmit-receive birdcage RF coil (Rapid Biomedical GmbH, Rimpar, Germany). Mice were anaesthetized with 1-3% isoflurane in 30% O_2_:70% N_2_ and positioned in the volume coil using a homebuilt cradle. Mouse temperature was monitored and maintained at 37 ± 0.5°C with a rectal probe and heating blanket feedback system^43^. Respiration was monitored using a pressure balloon.

A T_1_ weighted spin-echo multi-slice (SEMS) anatomical scan was acquired first with parameters: TR = 0.5 s, TE = 20 ms, number of averages = 2, matrix size = 256 x 256, FOV = 20 x 20 mm, in plane resolution = 78 μm isotropic, slice thickness 1 mm, number of slices = 8. A T_2_ weighted fast spin-echo multi-slice (fSEMS) anatomical scan was also acquired with TR = 3.5 s, echo spacing = 15 ms, effective TE = 60 ms, number of segments = 4, matrix size = 256 x 256, FOV = 20 x 20 mm, in plane resolution = 78 μm isotropic, slice thickness 1 mm, number of slices = 8.

A T_2_*-weighted three-dimensional multi-gradient echo (MGE3D) sequence was performed prior to contrast administration with parameters: repetition time (TR) = 32.55 ms, 1^st^ echo time (TE) = 2.5 ms, echo spacing (TE_2_) = 4 ms, number of echoes = 6, flip angle (FA) = 13°, matrix = 256 x 192 x 192 (zero-filled to 256 x 256 x 256), FOV = 22.5 x 22.5 x 22.5 mm, resolution = 88 μm isotropic. Subsequently, MPIO (4 mg Fe/kg in 100 μL sterile saline for all MPIO) were injected via a tail vein; 20 min later a postcontrast MGE3D scan was performed, as above, to detect MPIO binding.

Following this, 30 μL of gadodiamide (Omniscan, GE Healthcare, UK) was administered via the tail vein, and a post-gadolinium T_1_ weighted spin-echo multi-slice (SEMS) scan was acquired, as described previously, 5 min after gadolinium administration to assess BBB breakdown.

### MRI data analysis

The images were reconstructed for individual echoes of MGE3D using the square root of sum of squares (SqrtSOS) algorithm by an in-house MATLAB script. A segmentation mask for the striatum and any adjacent tumour bearing areas in the ipsilateral striatum, along with a contralateral striatum mask, was created for each mouse using ITK-SNAP software^44^. For the contralateral (normal) striatum, the mean signal intensity and standard deviation were calculated in MATLAB. Any voxel with a signal intensity of more than 3 standard deviations below the mean of the contralateral striatum was designated as hypointense. The total number of hypointense voxels in both the tumour-bearing and contralateral striatum was recorded. As the tumours contained naturally hypointense areas in the pre-MPIO images, all data are reported as the difference in hypointense voxels between the post- and pre-contrast scans (i.e. MPIO induced hypointense voxels). In a number of tumours from both targeted and control groups (n = 3 RGD-MPIO, n = 5 control RDG-MPIO), the hypointense voxels in the tumour-bearing striatum of the precontrast image appeared to be greater than those in the post-contrast image. As a result, an apparently negative difference in post-minus pre-contrast hypointense voxels was found. Images were inspected to ensure no artefacts were present that would affect image quality, or any differences in SNR. In the absence of any technical basis for this observation, it was determined that MPIO-induced hypointense voxels in these mice fall within the noise range of this technique (47 ± 31 voxels, ~0.2% of contralateral striatum volume). Consequently, for further analysis those animals were assigned a value of zero MPIO-induced hypointense voxels rather than a negative value, which is not possible.

### Immunohistochemistry

Brain tissue sections were stained for blood vessel marker CD31 (AF3628, R&D Systems, Abingdon, UK), VCAM-1 (1510-14, Southern Biotech, Birmingham, USA) and the macrophage/microglia marker Iba-1 (ab5076, Abcam, Cambridge, UK), as described previously^45–47^. Secondary antibodies used were biotinylated horse anti-goat IgG (BA-9500, Vector Laboratories, Peterborough, UK), and biotinylated goat anti-rat IgG (BA-9401, Vector Laboratories, Peterborough, UK). Sections were also stained for the angiogenic marker integrin α_v_β_3_ (ab78289, Abcam, UK) using the Mouse on Mouse Basic Kit (BMK-2202, Vector Laboratories, UK) in accordance with the protocol described by the manufacturer.

Microvessel density (CD31 positive area fraction) was quantified for the area encompassing the metastatic foci in the injected striatum, and also for the contralateral striatum, as the percentage of area covered using the “Positive Pixel Count 2004-08-11” algorithm in ImageScope (Leica Biosystems). The parameters used were moderately stained pixel intensity, between 202 and 185, and strongly stained pixel intensity, < 10.

### Perls’ Prussian blue staining

Perls’ Prussian blue staining was carried out on brain tissue sections to visualize ferric (+3) iron, including MPIO. Sections were washed in distilled water, then incubated in equal parts 20 mg/mL potassium ferrocyanide, and 2% v/v HCl in distilled water for 15 minutes. Sections were then washed in distilled water, followed by counterstaining with nuclear fast red for 2 minutes. Sections were then dehydrated and mounted with coverslips.

Where double staining for molecular markers and Perls’ Prussian blue was performed, the immunohistochemistry protocol described above was carried out to the point of visualization with DAB, and the Prussian blue staining protocol then applied as above. Following Prussian blue staining, sections were either counterstained with nuclear fast red (Iba-1, VCAM-1) or proceeded directly to dehydration and coverslip mounting (CD31).

### Assessment of in vitro uptake of MPIO

4T1-GFP tumour cells and RAW-264.7 murine macrophages^48^ were seeded onto 6-well plates in Dulbecco’s Modified Eagle’s Medium (DMEM) supplemented with 1% L-glutamine and 10% (v/v) fetal bovine serum (FBS). When cells reached approximately 80% confluency, 1 μg of RGD-MPIO, control RDG-MPIO, VCAM-MPIO or IgG MPIO were added to separate wells. Cells were incubated in a shaking incubator for 40 minutes, then washed 5 times with PBS to remove any unbound MPIO. Cells were fixed with 4% paraformaldehyde and then examined under a Nikon Ti-E inverted microscope. Images were taken at random locations in each well, and MPIO were quantified by manual counting.

To analyse receptor-specific uptake in RAW264.7 macrophages, cells were seeded onto 6-well plates, as above. Cells were pre-treated for 1 hour with 0.1, 1 or 10 μL of 10 mM RGD or control RDG peptide; 0.1, 1 or 10 μg VCAM-1 or IgG antibody. Following this pre-treatment, 1 μg of the same MPIO as the blocking agent was administered. Cells were incubated, washed, fixed and examined by microscope, as above.

### Endogenous iron-laden macrophages

In order to determine whether endogenous iron-laden macrophages were present in the 4T1-GFP brain macrometastasis model across the tumour timecourse, additional cohorts of mice were sacrificed at day 7 (n = 4), 14 (n = 4), 21 (n = 2), 28 (n = 4) or 35 (n = 3) and the brains taken for immunohistochemical analysis. These animals had no prior exposure to iron oxide contrast agents.

A further cohort of animals underwent MRI at a single timepoint, day 28 (n = 6) to determine whether endogenous iron in tumour-associated macrophages produces non-specific hypointense voxels in precontrast images. A T_2_*-weighted MGE3D dataset was acquired, but no MPIO administered, and a gadolinium-enhanced T_1_ weighted scan was also performed. Either single staining with Prussian blue or double staining for Iba-1 and Prussian blue was carried out on brain tissue sections, as described above, and iron-laden macrophages manually counted.

### Statistical analysis

Statistical analysis was carried out using Prism version 8.1.2. Normally distributed groupwise data were analysed using one-way ANOVA with Holm-Sidak’s post-hoc multiple comparison tests. Where only two groups exist, an unpaired student’s t-test with Welch’s correction was used.

Groupwise data which were not normally distributed were analysed using a Kruskal-Wallis one-way ANOVA with Dunn’s post-hoc multiple comparison tests. Paired groupwise data (i.e. comparing ipsilateral and contralateral values across timepoints) were analysed using two-way repeated measures ANOVA with Sidak’s multiple comparison tests.

Correlations were analysed using the Spearman r test. Linear regression lines were compared using the sum-of-squares F test.

## Results

### In vitro binding of cyclic RGD peptides to mouse integrin α_v_β_3_

To determine the binding efficacy of various RGD peptides to the mouse integrin α_v_β_3_, the number of RGD-MPIO binding events in α_v_β_3_ or control (BSA coated) glass capillaries was compared under flow conditions (Supplementary Figure 2). Comparison of binding events between α_v_β_3_ and BSA capillaries showed an effect of both capillary (2-way ANOVA, p < 0.0001) and cyclic-RGD (2-way ANOVA, p < 0.001). Tukey’s multiple comparison post-hoc tests showed significant differences between BSA and α_v_β_3_ coated capillaries for the RGD-1 (p < 0.05), RGD-2 (p < 0.001) and RGD-3 (p < 0.001) cyclic-RGD peptides (Supplementary Figure 2). The control RDG peptide showed no significant difference in binding to BSA or α_v_β_3_ coated capillaries (Supplementary Figure 2). For the α_v_β_3_ capillary, the RGD-2 peptide showed significantly higher binding than both the control RDG peptide (1 way ANOVA p < 0.005, Bonferroni’s multiple comparison post-hoc test p < 0.001), and the other RGD peptides RGD-1 (post-hoc test p < 0.001) and RGD-3 (post-hoc test p < 0.05). Thus, peptide RGD-2 was used for further experiments with RGD-MPIO, and peptide RDG was used as a negative control scrambled peptide for RDG-MPIO.

In blocking experiments, to assess the specificity of RGD-MPIO binding to α_v_β_3_ on endothelial cells in culture (Supplementary Figure 3), significantly higher binding was observed for RGD-MPIO to SNAP stimulated HUVEC cells than either unstimulated cells (Welch’s ANOVA with Dunnett’s multiple comparison test, p < 0.001) or cells pre-treated with an RGD block (p < 0.01). No differences were found for control RDG-MPIO binding to stimulated cells, with or without an RGD block, compared to unstimulated cells. Together the above findings demonstrate specificity of RGD-MPIO binding to α_v_β_3_.

### Assessment of angiogenesis and α_v_β_3_ expression in a mouse model of brain macrometastasis

For assessment of vessel density, correlation analysis showed that both the CD31 stained area and number of CD31 positive vessels increased more rapidly over time in the tumour-bearing striatum than in the contralateral striatum (sum-of-squares F test, p < 0.05 for both; Figure 1A-B). No overlap between tumour and contralateral striatum regression 95% confidence intervals was evident past day 12 or 13, respectively, indicating an increase in tumour blood vessels (i.e. angiogenesis) after this point.

**Figure 1.**
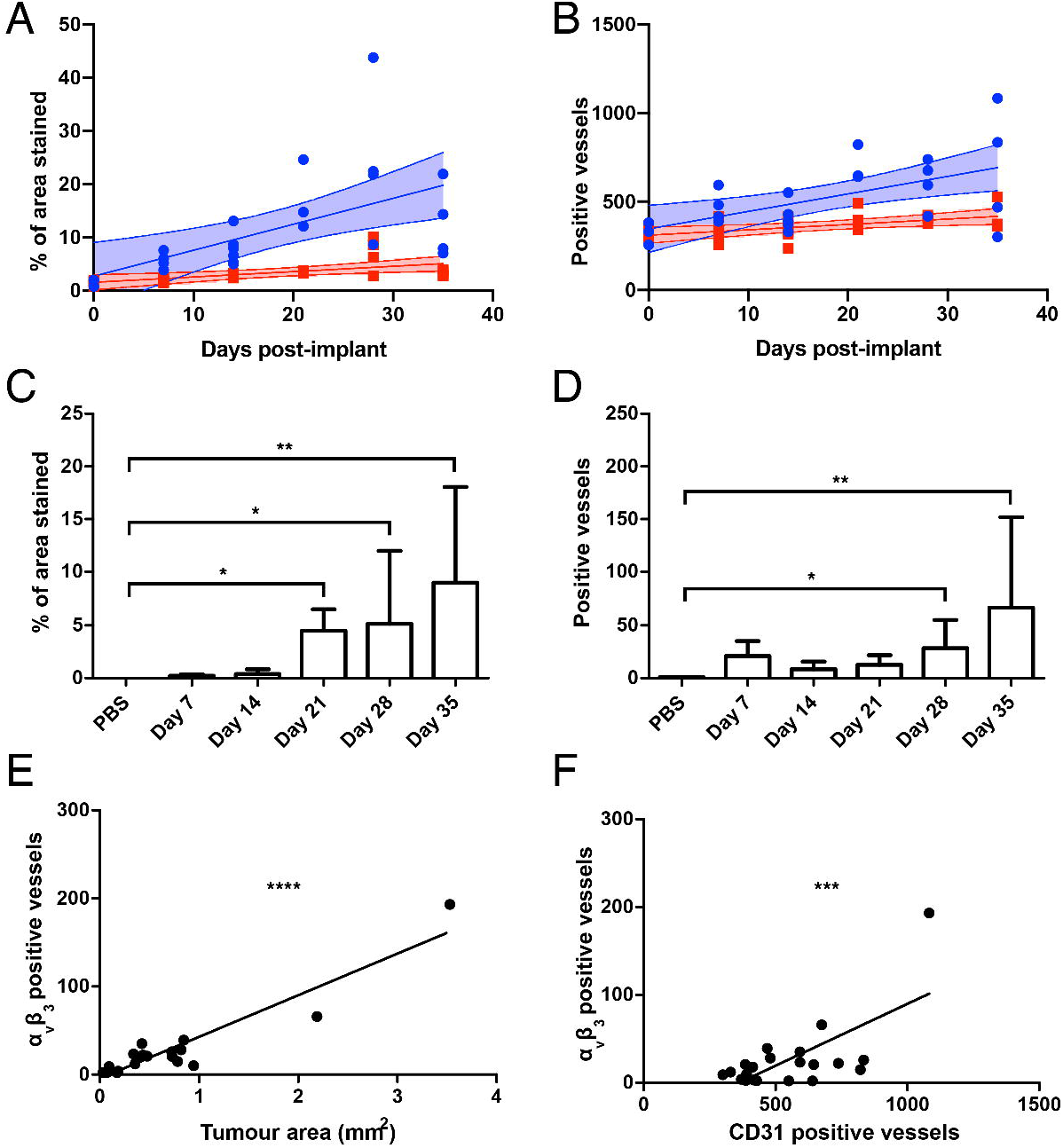
(A) Quantitation of CD31 total stained area as a percentage of tumour area in the tumour (blue circles) and the contralateral striatum (red squares), plotted against time with 95% confidence intervals. A significant increase in blood vessels in the tumour is evident after day 12 (linear regression sum-of-squares F-test p < 0.05; Spearman’s r, tumour regression: 0.75, p < 0.0001; contralateral striatum regression: 0.74, p < 0.0001). (B) Comparison of microvessel density (CD31 positive vessels) in the tumour (blue circles) and the contralateral striatum (red squares), plotted against time with 95% confidence intervals. A significant increase in blood vessels in the tumour is evident after day 13 (linear regression sum-of-squares F-test p < 0.05; Spearman’s r, tumour regression: 0.56, p < 0.01; contralateral striatum: 0.51, p<0.05). (C) Quantitation of total α_v_β_3_ stained area as a percentage of tumour area across the timecourse. (D) Quantitation of α_v_β_3_ positive vessels across the timecourse. (E) Number of α_v_β_3_ positive vessels correlated positively with tumour size (Pearson’s coefficient p < 0.0001, R^2^ = 0.88). (F) Number of α_v_β_3_ positive vessels correlated positively with CD31 microvessel density (Pearson’s coefficient p < 0.001, R^2^ = 0.46). In all cases, PBS control (n = 3), day 7 (n = 4), 14 (n = 6), 21 (n = 3), 28 (n = 4), and 35 (n = 4). Data for these figures had a lognormal distribution, and no outliers were found to be present (ROUT test). Bars represent mean ± standard deviation; *p < 0.05, **p < 0.01, ***p < 0.001.

Immunohistochemical staining revealed a significant increase in α_v_β_3_ total stained area as a percentage of tumour area, compared to control PBS injected animals, from day 21 (Kruskal-Wallis test p < 0.01, post-hoc p < 0.05 (day 21,28) or p < 0.01 (day 35); Figure 1C). Further, a significant increase in α_v_β_3_ positive vessels, specifically, compared to control PBS injected animals, was evident at days 28 and 35 (Kruskal-Wallis test p < 0.05, post-hoc p < 0.05 (day 28) or p < 0.01 (day 35); Figure 1D). Significant positive correlations were found between α_v_β_3_ positive vessels and both tumour area (p < 0.0001, Pearson’s coefficient R^2^ = 0.88; Figure 1E) and CD31 positive vessels (p < 0.001, Pearson’s coefficient R^2^ = 0.46; Figure 1F).

### Imaging angiogenesis in vivo using RGD-MPIO

Having demonstrated the presence of α_v_β_3_ positive angiogenic vessels in this model of brain macrometastasis, the ability of the chosen RGD-MPIO to detect these vessels *in vivo* was assessed across a timecourse. Qualitatively, increasing numbers of hypointensities were apparent in the tumour-bearing striatum on T_2_*-weighted images following intravenous RGD-MPIO injection across the timecourse studied (Supplementary Figure 4), with 80% of mice showing MPIO-induced hypointensities by day 35 (Figure 2Ai-ii). In contrast, few hypointensities were observed in mice injected with RDG-MPIO at the early timepoints (days 7-21, Supplementary Figure 5), but became more evident at days 28 and 35 (57% of mice; Figure 2Aiv and Supplementary Figure 5). Some hypointense voxels were also apparent in the pre-contrast T_2_*-weighted MGE3D images in the tumour bearing striatum (Figure 2A and Supplementary Figures 4 and 5). Consequently, all following MPIO induced hypointensity values are reported as post-minus pre-contrast hypointensities.

**Figure 2.**
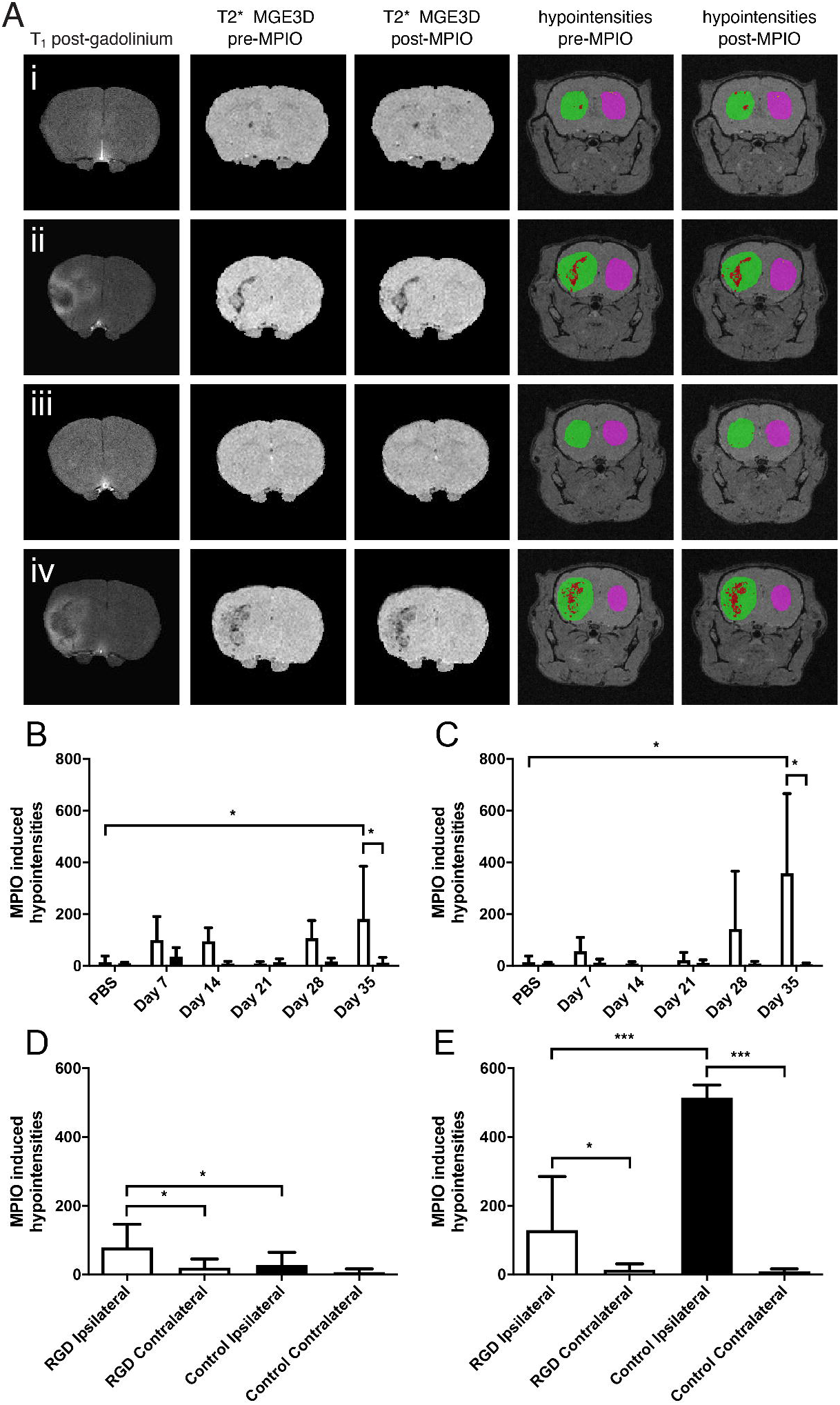
(A) Representative MR images from mice bearing 4T1-GFP macrometastases at day 28-35. (Ai,ii) Tumours imaged with RGD-MPIO; (Ai) non-enhancing tumour at day 28, (Aii) gadolinium-enhancing tumour at day 35. (Aiii-iv) Tumours imaged with RDG-MPIO; (Aiii) non-enhancing tumour at day 28, (Aiv) gadolinium-enhancing tumour at day 35. Each column of images, from the left, shows T_1_-weighted post-gadolinium images, T_2_*-weighted pre-MPIO MGE3D images, T_2_*-weighted post-MPIO MGE3D images, and overlays on T_2_*-weighted MGE3D images showing hypointensities pre-MPIO and post-MPIO. For the overlays, the tumour-bearing striatum is segmented in green, and the contralateral striatum segmented in pink. Hypointense voxels are shown in red. (B) Significantly increased RGD-MPIO induced hypointense voxels were evident in the tumour-bearing striatum (white bars) compared to the contralateral striatum (black bars) at day 35 (2-way paired ANOVA p < 0.05). (C) Significantly increased control RDG-MPIO induced hypointense voxels were also seen in the tumour-bearing striatum (white bars) compared to the contralateral striatum (black bars) at day 35 (2-way paired ANOVA p < 0.05). (DE) Comparison of pooled data across all timepoints for (D) non-enhancing tumours, and (E) gadolinium-enhancing tumours. (D) Mice with non-enhancing tumours administered RGD-MPIO (white bars; n=7) showed significantly increased MPIO-induced hypointense voxels in the tumour-bearing hemisphere (1-way ANOVA p < 0.005) than both the contralateral hemisphere and the mice administered control RDG-MPIO (black bars; n=13) in the tumour-bearing hemisphere. (E) In mice with gadolinium-enhancing tumours, significantly increased MPIO-induced hypointense voxels were observed in the tumour-bearing striatum compared to the contralateral striatum for both RGD-MPIO (white bars; n=10) and RDG-MPIO (black bars; n=3) (1-way ANOVA p < 0.0001). However, in mice receiving control RDG-MPIO, the volume of MPIO-induced hypointense voxels was also significantly greater than those administered RGD-MPIO. Number of MPIO induced hypointense voxels, presented as post-contrast minus pre-contrast hypointense voxels for all data. Bars represent mean ± standard deviation; post-hoc Holm-Sidak’s tests *p < 0.05, **p < 0.01, ***p < 0.001.

Quantitative analysis of MPIO induced hypointensities in mice injected with RGD-MPIO showed significantly more in the tumour-bearing *vs.* contralateral striatum at day 35 (2-way paired ANOVA p < 0.05, post-hoc Sidak’s test p < 0.05; Figure 2B). However, in mice injected with the control RDG-MPIO, a significant difference was also observed between the tumour-bearing and contralateral striatum at day 35 (2-way paired ANOVA p < 0.05, post-hoc Sidak’s test p < 0.05; Figure 2C). In mice receiving the PBS control injection (no tumour), very few hypointense voxels were detected in mice receiving MPIO in either the ipsilateral or the contralateral striatum (Figure 2B-C). At face value, these findings suggest that nonspecific retention of MPIO occurred in tumour-bearing mice. However, tumour growth and progression were found to be variable over the timecourse, with some tumours at the later timepoints showing marked BBB breakdown and gadolinium enhancement (Figure 2Aii, iv), whilst others had an intact BBB (Figure 2Ai, iii). Consequently, to determine why significant uptake of control RDG-MPIO was observed at the day 35 timepoint, gadolinium-enhancing and non-enhancing tumours were separated and pooled across timepoints for further analysis (Supplementary Table 1).

In the pooled group of non-enhancing tumours (n = 7 RGD-MPIO, n = 13 control RDG-MPIO), mice injected with RGD-MPIO showed significantly increased MPIO-induced hypointense voxels in the tumour-bearing hemisphere (1-way ANOVA p < 0.005; Figure 2D) than both the contralateral hemisphere (Holm-Sidak’s post-hoc test p < 0.05), and the mice administered control RDG-MPIO in the tumour-bearing hemisphere (Holm-Sidak’s post-hoc test p < 0.05). No difference was observed between the tumour-bearing and contralateral striatum in mice administered control RDG-MPIO (Figure 2D). In contrast, for the gadolinium-enhancing tumours (n = 10 RGD-MPIO, n = 3 control RDG-MPIO, from days 21, 28 and 35), mice administered with either RGD-MPIO or control RDG-MPIO both showed significantly more MPIO-induced hypointense voxels in the tumour-bearing striatum than the contralateral striatum (1-way ANOVA p < 0.0001, Holm-Sidak’s post-hoc test p < 0.05 RGD-MPIO or p < 0.001 RDG-MPIO; Figure 2E). Moreover, the number of hypointense voxels in the control RDG-MPIO group was greater than the group given RGD-MPIO (Holm-Sidak’s post-hoc test p < 0.001). In both groups the number of hypointensities observed was considerably greater than for the non-enhancing tumours.

### Non-specific tumour retention of MPIO by macrophages

The above data raised the question of whether peptide-targeted MPIO are taken up non-specifically within the tumour microenvironment. Tissue sections from mice with gadolinium-enhancing tumours were stained with Perls’ Prussian blue to visualize iron in the tumour tissue. Whilst individual MPIO were visible in the tumour tissue associated with the vascular endothelium (Figure 3A, black arrowheads), large blue areas indicating aggregations of iron were also visible in tissue sections at all timepoints (Figure 3A, red arrowheads). These aggregations were observed in mice injected with both RGD-MPIO and control RDG-MPIO, across all timepoints. A significant positive correlation was observed between tumour size and number of aggregations in mice given RGD-MPIO (p < 0.0005, Pearson’s coefficient R^2^ = 0.64; Figure 3B), but not control RDG-MPIO (Figure 3C), although a similar trend was apparent.

**Figure 3.**
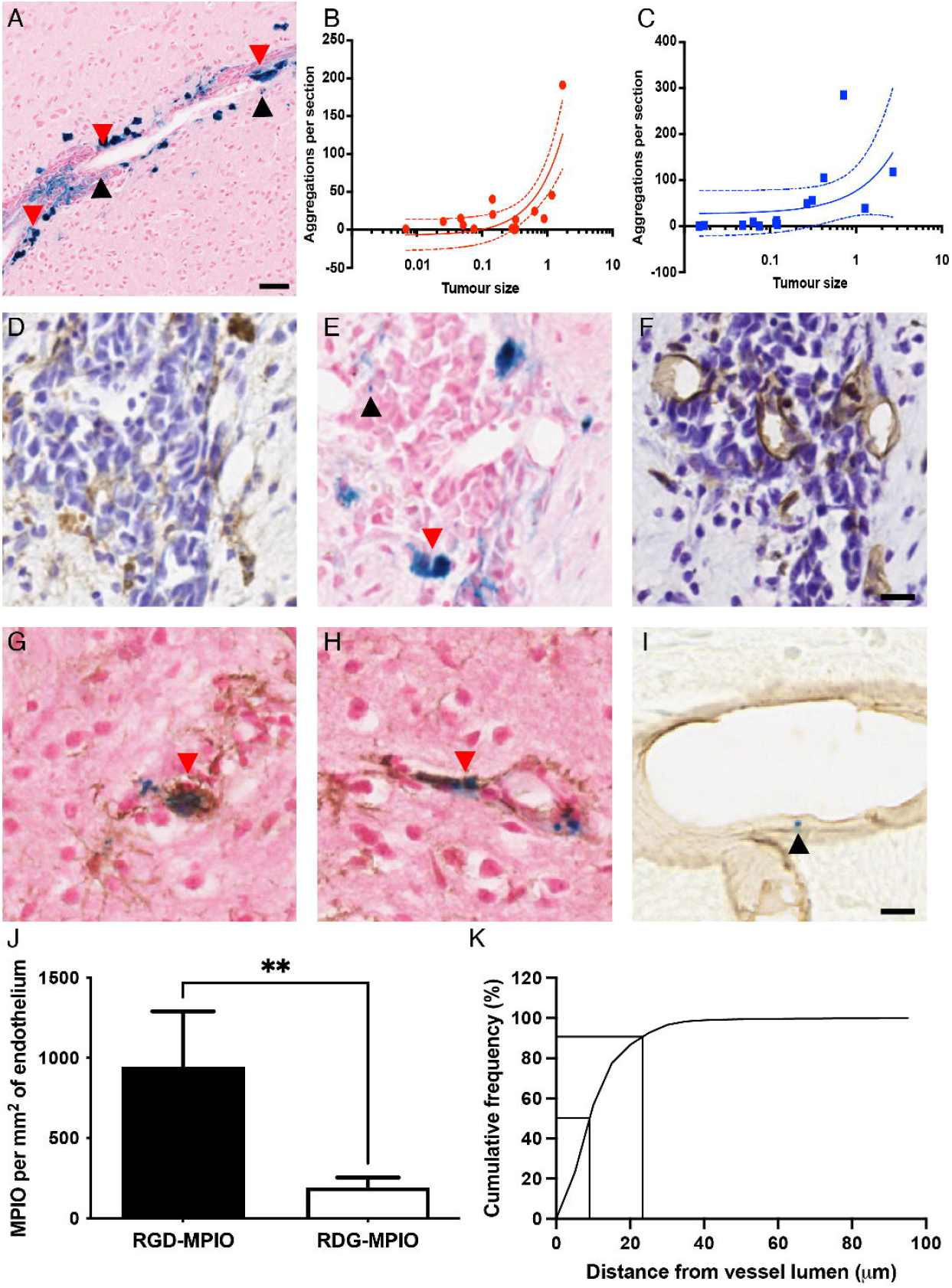
Aggregations of iron in 4T1-GFP tumour tissue. (A) Representative section of gadolinium enhancing tumour stained for Perls’ Prussian blue (iron, blue) and counterstained with nuclear fast red. Black arrowheads indicate examples of single MPIO associated with the vascular endothelium, and red arrowheads indicate examples of iron aggregations. (B-C) Correlations between iron aggregations and tumour size in mice injected with either (B) RGD-MPIO or (C) control RDG-MPIO. Linear regression analysis showed a positive correlation between number of aggregations and tumour size (R^2^ = 0.64, ***p < 0.001) in mice injected with RGD-MPIO, but not control RDG-MPIO, although a similar trend was evident; 95% confidence intervals shown. (D-F) Consecutive sections stained for (D) Iba-1 (macrophages and microglia, brown staining), (E) Prussian blue (iron) and (F) CD31 (blood vessels, brown staining) in a day 35 mouse injected with RGD-MPIO. Red arrow indicates a larger iron aggregation in similar location to macrophage staining (D; Iba-1), whilst black arrow indicates single MPIO distant from macrophage staining, but close alignment with a blood vessel (F; CD31). (G-H) Double staining of 4T1-GFP tumour tissue sections, from a mouse injected with RGD-MPIO at the day 35 timepoint, for Prussian blue (iron) and macrophages/microglia (Iba-1, brown staining), indicate co-localisation of iron within macrophages/microglia (red arrows). Sections counterstained with nuclear fast red. (I) Double staining for Prussian Blue and CD31 revealed single MPIO (black arrow) bound to blood vessels (brown stained) in mice injected with RGD-MPIO; representative image from day 21 shown. (J) In the gadolinium enhancing 4T1-GFP tumours, single endothelium bound MPIO are observed more often in mice injected with RGD-MPIO (n = 10) than control RDG-MPIO (n = 3, t-test, **p < 0.01). (K) Histogram showing the cumulative frequency of the measured distance between the centre of the iron-laden macrophages and the centre of the blood vessel lumen, indicating their close association with blood vessels. Scale bar = 25 μm in (A) and 10 μm in (D-I).

Staining of consecutive tumour sections revealed that the iron aggregations were located in similar spatial patterns to macrophages (Figure 3D-E). In contrast, single MPIO could be clearly identified in regions distant from macrophage staining and in close association with vessels (Figure 3E-F). Double staining of Perls’ Prussian blue with the macrophage/microglia marker Iba-1 confirmed that the aggregations of iron were present inside macrophages/microglia (Figure 3G-H). In contrast, double staining of Perls’ Prussian blue with the endothelial marker CD31 revealed single MPIO bound to vessels (Figure 3I).

Quantitation of single MPIO associated with blood vessels showed significantly more RGD-MPIO bound per mm of endothelium than control RDG-MPIO (t-test, p < 0.01; Figure 3J). Double staining for CD31 and Prussian blue indicated that over 90% of iron-laden macrophages were within 25 μm of the blood vessel lumen, with a median distance of 10 μm (Figure 3K), suggesting that they are likely to be perivascular macrophages.

### Effect of MPIO targeting moiety

Since retention of MPIO by perivascular macrophages has not previously been observed in studies using antibody-targeted MPIO to study brain pathology^33,47,49–51^, we next sought to determine whether peptide-targeted MPIO are taken up by perivascular macrophages preferentially and could explain, at least in part, the observed iron aggregations in the current study.

Firstly, *in vitro* uptake of both peptide- and antibody-targeted MPIO by cultured macrophages was assessed. Substantial numbers of MPIO were taken up by RAW 264.7 macrophages *in vitro* for all four MPIO types tested (Supplementary Figure 6A), whilst significantly fewer MPIO were taken up by cultured 4T1-GFP cells (Supplementary Figure 6B; 2-way ANOVA, p < 0.001). However, no significant differences in uptake by RAW 264.7 macrophages were observed following 1 hour of blocking with either RGD or control RDG peptides for RGD-MPIO and RDG-MPIO, respectively (Supplementary Figure 6C-D), or after 1 hour of blocking with VCAM-1 or IgG antibodies for VCAM-MPIO and IgG-MPIO, respectively (Supplementary Figure 6E-F). These results indicate that uptake by macrophages in culture is neither receptor nor ligand-type dependent.

Next, accumulation of antibody-conjugated MPIO in perivascular macrophages *in vivo* was assessed in 4T1-GFP macrometastases at day 28 (Figure 4A). Mice were injected with either targeted VCAM-MPIO or control IgG-MPIO (Figure 4B-C). VCAM-1 has previously been shown to be upregulated on the vascular endothelium in close association with brain metastases^47,49,52,53^. In the mice receiving VCAM-MPIO, MPIO induced hypointense voxels were observed in the tumour-bearing, but not the contralateral, striatum (Figure 4D). Within this cohort of mice a significant variation in tumour size was evident, with one animal exhibiting a relatively small, non-enhancing tumour (0.5 mm^3^) whilst the other two had much larger, contrast-enhancing tumours (5.5 and 18.2 mm^3^, respectively). In the mouse with the smaller tumour, only a relatively small number of hypointense voxels was detected, resulting in a large spread in effect size across the group (Figure 4D) and, consequently, the difference between hemispheres did not reach statistical significance. Nevertheless, it is clear that all of the animals injected with VCAM-MPIO showed more hypointense voxels in the tumour bearing than contralateral hemisphere. Numbers of hypointense voxels were considerably higher than observed following RGD-MPIO administration. In mice receiving the control IgG-MPIO, no MPIO induced hypointense voxels were observed in either striatum, despite the presence of large (2.9 and 8.6 mm^3^, respectively) contrast-enhancing tumours (Figure 4D).

**Figure 4.**
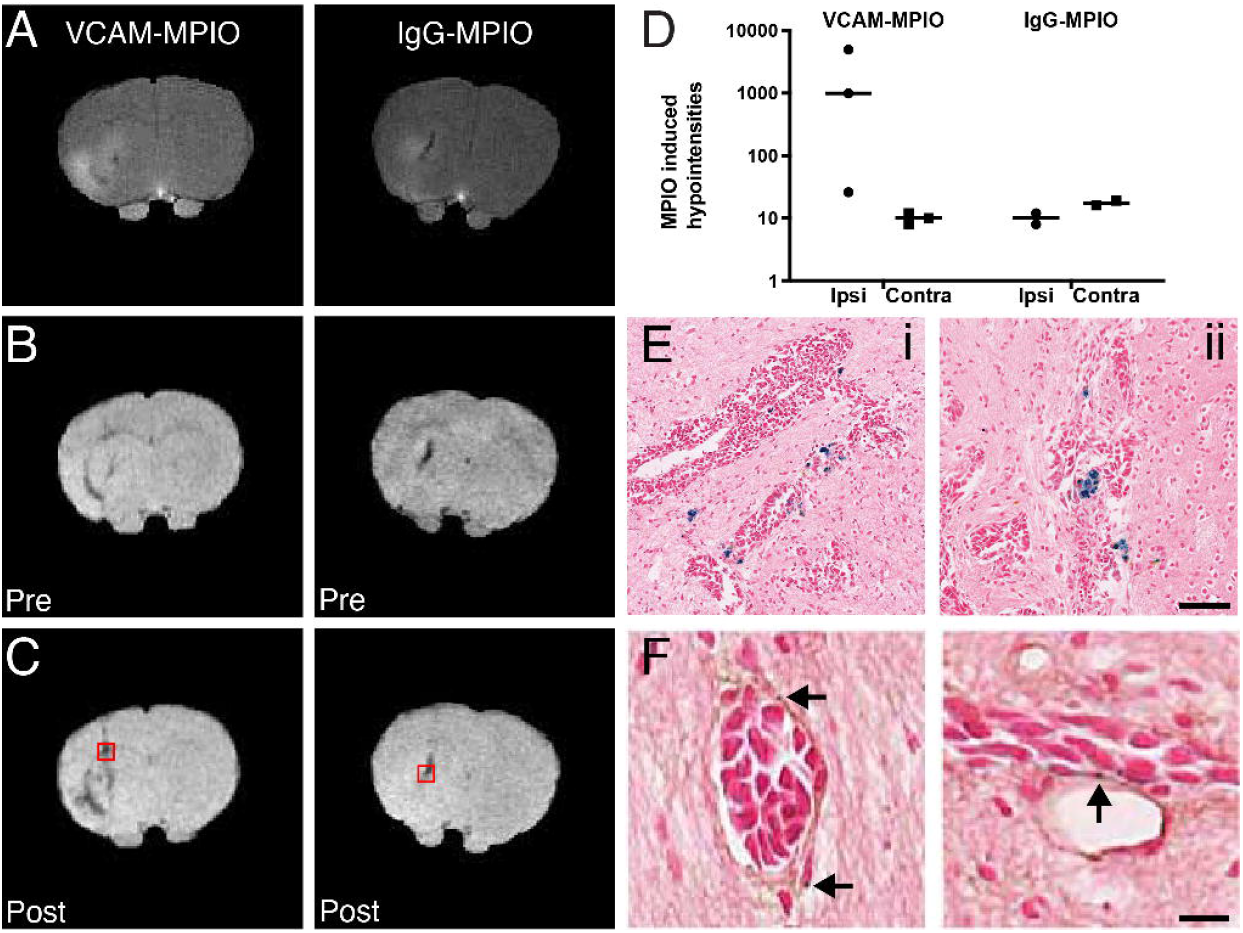
MR and histology images from mice with 4T1-GFP brain metastases. (A) Representative postgadolinium T_1_-weighted images from two different mice, injected with either VCAM-MPIO or IgG-MPIO, demonstrating contrast enhancement in the tumour-bearing striatum. (B-C) Representative pre-MPIO (B) and post-MPIO (C) MGE3D images in mice imaged with either VCAM-MPIO or IgG-MPIO. (D) Quantitation of MPIO induced hypointense voxels, presented as post-contrast minus pre-contrast for ipsi- and contralateral hemispheres in each individual animal. Bars indicate mean. (E) Histology images showing Prussian blue staining from the same animals post-MRI, and corresponding to the regions in red squares in (C), respectively; the location of apparent MPIO-laden macrophages corresponds to hypointense regions in the pre-contrast MGE3D images (B). Scale bar = 100 μm. (F) In mice imaged with VCAM-MPIO, specific endothelial binding of individual VCAM-MPIO (arrows) to VCAM-1 positive blood vessels (brown) can be seen. Scale bar = 10 μm.

Analysis of Prussian blue histological staining from animals injected with antibody-targeted MPIO (Figure 4E) showed similar levels of iron-laden macrophages between mice injected with VCAM-MPIO (Figure 4Ei) or IgG-MPIO (Figure 4Eii, Supplementary Figure 7, 40.9 ± 52.4 *vs.* 50.2 ± 64.8 per tissue section), with no significant differences between groups. These numbers of iron-laden macrophages were also not significantly different to those observed at the same time-point in mice injected with peptide-targeted MPIO (17.8 ± 16.2 per tissue section; Supplementary Figure 7). Moreover, further analysis indicated that all such macrophages were located in areas exhibiting hypointensity on T_2_*-weighted images prior to MPIO injection (Figure 4B). This observation, together with the lack of MPIO-induced hypointensities observed in IgG-MPIO injected mice, suggests that these iron-laden macrophages do not contain MPIO, but rather endogenous iron. Double staining for Prussian blue and VCAM-1 revealed that in addition to iron-laden macrophages, single MPIO were present bound to VCAM-1 positive blood vessels in mice imaged with VCAM-MPIO (Figure 4F), but not in mice imaged with control IgG-MPIO.

### Endogenous iron-laden macrophages

The above findings led us to investigate whether the larger iron aggregations observed were indeed endogenous iron rather than MPIO, since they correlated spatially with pre-existing hypointensities. Thus, mice bearing 4T1-GFP tumours, but with no exposure to iron oxide contrast agents, were assessed for the presence of endogenous iron-laden macrophages. Low levels of endogenous iron-laden macrophages were observed in the tumour tissue across all timepoints (Figure 5A-B and Supplementary Figure 7) and were histologically indistinguishable from the iron-laden macrophages observed in the previous studies, no statistically significant differences were evident between time points. When endogenous iron-laden macrophage levels were compared to tumour size, no significant correlation was found (p > 0.2, Pearson’s coefficient R^2^ = 0.08; Supplementary Figure 8A).

**Figure 5.**
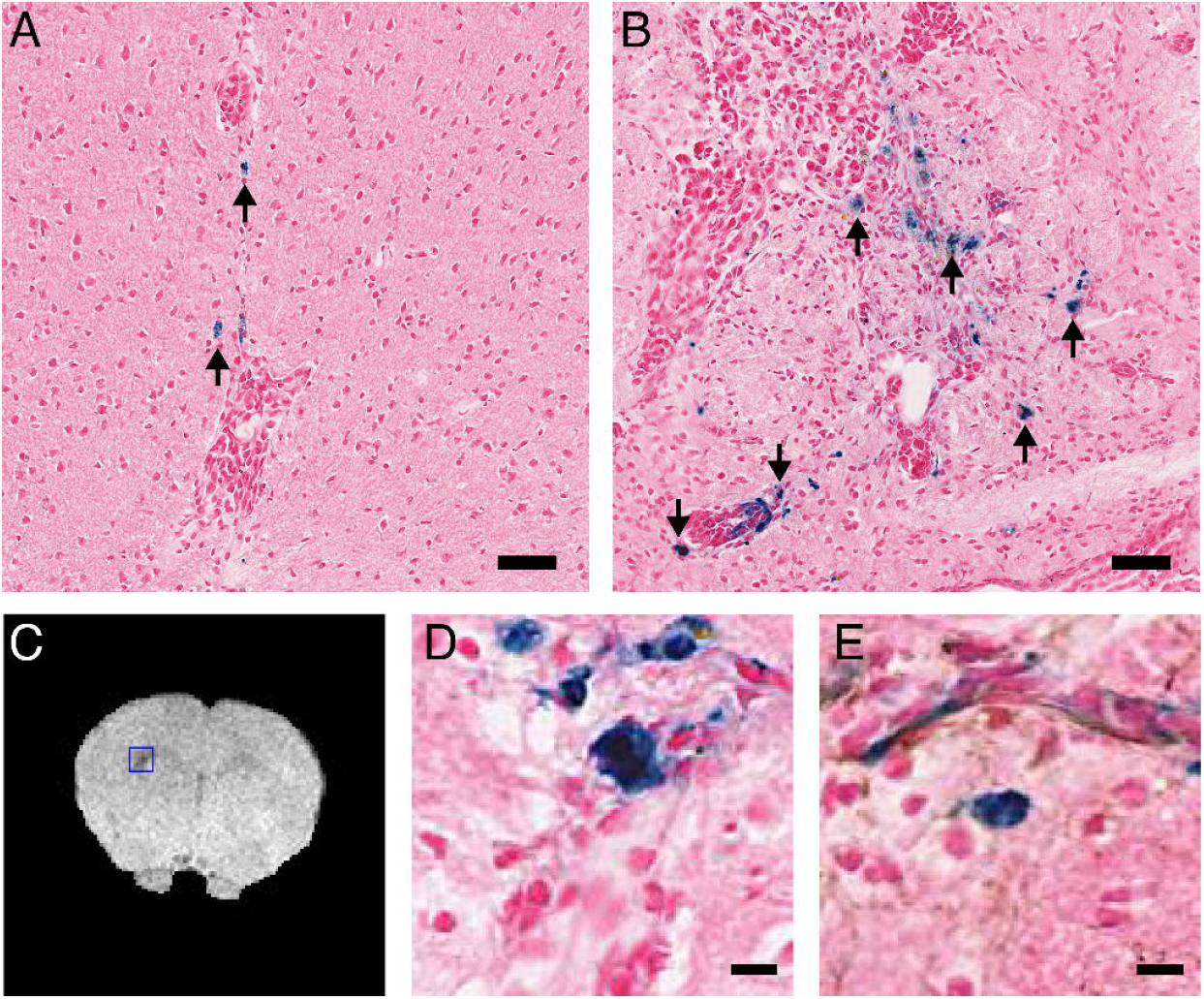
Endogenous iron-laden macrophages in metastases with no exposure to MPIO correspond to pre-existing hypointensities. (A-B) Histology sections stained for Prussian blue (iron, blue), and counterstained with nuclear fast red (nuclei, pink). Arrows indicate iron-laden macrophages. Scale bar = 40 μm. Representative images from (A) day 7, and (B) day 28. (C) Hypointense voxels are evident in the ipsilateral striatum on T_2_*-weighted images (blue box). (H-I) Histologically, the location of endogenous iron-laden macrophages corresponds to hypointense voxels in the MGE3D images; sections stained for (H) iron (Prussian blue) or (I) double stained for iron and macrophages/microglia (Iba1, brown) shown at 200x magnification for the areas indicated by blue box in the MR image. Sections are counterstained with nuclear fast red (nuclei, pink). Scale bars indicate 50 μm in (A-B) and 10 μm in (D-E).

T_2_*-weighted images from 4T1-GFP tumours at the day 28 timepoint (Figure 5C) were compared with Prussian blue stained histological sections. The tumours showed varying levels of pre-existing hypointense voxels in the tumour bearing striatum (720 ± 570; Supplementary Figure 8B). In contrast, the contralateral striatum exhibited very few pre-existing hypointense voxels (57 ± 21; Supplementary Figure 8B), and a significant difference was evident between hemispheres (paired t-test, p < 0.05). The levels of pre-existing hypointense voxels in this study are similar to the levels of pre-contrast hypointense voxels observed in the 4T1-GFP bearing animals subsequently imaged with RGD-MPIO or control RDG-MPIO. Spatially, the endogenous iron-laden macrophages correlated closely with pre-existing hypointense voxels in the T_2_*-weighted images (Figure 5C-D). Double staining for Prussian blue and the macrophage/microglia marker Iba-1 showed co-localisation of iron aggregations and Iba-1 staining (Figure 5E). These data suggest that there is a baseline level of endogenous iron-laden macrophages that underlie the pre-contrast hypointensities observed in the above studies.

### MPIO penetration of the blood-brain barrier

Given that the numbers of iron-laden macrophages did not appear to differ significantly between groups, irrespective of targeting moiety or indeed presence of MPIO, we next tested the hypothesis that peptide-targeted MPIO are able to simply extravasate across the BBB when frank breakdown is evident. To this end, a CINC-1 induced model of BBB breakdown was used. As non-specific accumulation of MPIO had not previously been observed in models of brain macrometastasis when targeted with antibodies rather than peptides^52^, both peptide- and antibody-targeted MPIO were again evaluated. BBB breakdown was confirmed in all animals using gadolinium-enhanced imaging (Supplementary Figure 9A).

In mice across all groups, a small number of hypointense voxels post-MPIO injection were evident in the CINC-1 injected striatum, but not in the contralateral striatum (Supplementary Figure 9B,C), although the difference between hemispheres did not reach statistical significance. Histological examination of brain tissue, however, revealed only occasional single MPIO and a small number (fewer than 2 per section) of iron-laden macrophages (Supplementary Figure 6D-E). No differences were found between the different MPIO used. These data suggest that neither peptide- nor antibody-targeted MPIO cross a permeable BBB *per se* to any significant degree.

## Discussion

In this study, we have demonstrated that expression of the integrin α_v_β_3_ increases over time in this model of brain macrometastasis, and appears to correlate with increased vascularity and vessel permeability. These data suggest that integrin α_v_β_3_ has potential as a biomarker of angiogenesis in brain macrometastasis, and led to testing of the α_v_β_3_-targeted RGD-MPIO contrast agent. However, subsequent findings revealed a number of confounds that may reduce the sensitivity for detection of α_v_β_3_-specific binding using this peptide-targeted MPIO, including (i) the presence of endogenous iron-laden macrophages, particularly in larger brain metastases, which yield significant baseline hypointensities on T_2_*-weighted images, and (ii) non-specific retention of peptide-targeted MPIO within the tumour microenvironment. In contrast, antibody conjugated MPIO, despite specific binding to their endothelial target VCAM-1, showed negligible *in vivo* uptake by tumour macrophages and we propose, therefore, that the use of antibodies rather than peptides as the targeting ligand may represent a preferable route forward for new angiogenesis-targeted agents.

This study was a proof-of-concept study aiming to determine whether RGD-MPIO could detect expression of integrin α_v_β_3_ associated with tumour angiogenesis *in vivo.* In the model of directly induced 4T1-GFP brain macrometastasis used, endothelial α_v_β_3_ expression was found across the timecourse studied, but increased significantly at later timepoints together with increased microvessel density. Consequently, this was considered to be a suitable model to assess imaging of α_v_β_3_ as a biomarker of tumour angiogenesis in brain macrometastasis. Increased microvessel density and integrin α_v_β_3_ expression are both features of human breast cancer brain metastasis. It has been shown in both clinical samples^54^ and preclinical studies^55^ that breast cancer brain metastases have a higher microvessel density than their respective primary tumours. Breast cancer brain metastases have also been shown to express integrin α_v_β_3_^56,57^, so it is likely that if a successful integrin α_v_β_3_ imaging agent is developed it will be clinically relevant.

In our study, T_2_*-weighted imaging showed that pre-existing hypointensities could be observed prior to MPIO injection. Subsequent investigation of mice with tumours, but no exposure to iron oxide contrast agents, revealed endogenous iron-laden macrophages were present in the tumour tissue throughout the tumour timecourse, which correlated spatially with the pre-existing MRI hypointensities. This finding is supported by the results of Leftin *et al,* who have previously used T_2_* weighted imaging to quantify macrophages in a mouse model of brain metastasis^58^. Although the presence of iron-laden macrophages has not been observed in the micrometastatic stages of tumour development in the brain (<500 μm diameter), it is likely that endogenous iron-laden macrophages will be present at some level in most, if not all, brain tumours once they are more established. Thus, for any study using iron oxide-based contrast agents, it will be important to quantify any pre-existing hypointense voxels prior to administration of contrast.

Nevertheless, despite the presence of natural hypointensities, specific α_v_β_3_-targeted binding and MPIO induced hypointensities could be observed with RGD-MPIO, when only non-enhancing tumours were considered, with no retention of the control RDG-MPIO. Thus, whilst the background endogenous iron may reduce the sensitivity of this method, it does not necessarily preclude it and imaging of α_v_β_3_ positive angiogenic vessels may be possible in brain tumours if confounding factors can be eliminated. Moreover, this approach may be combined with complementary methods, such as arterial spin labelling (ASL) and dynamic contrast-enhanced MRI, to provide additional functional information on angiogenesis^59^.

In gadolinium enhancing tumours, however, retention of both the RGD-MPIO and the control RDG-MPIO was observed, resulting in large volumes of MPIO-induced hypointensities. This was an unexpected result, as the control RDG-MPIO had shown only very low levels of binding *in vitro* both here and in a previous study^35^. These data appeared to suggest that the MPIO, irrespective of targeting peptide, were non-specifically retained within the brain in the presence of frank BBB breakdown (as detected by gadolinium enhancement). Analysis of the histology from these animals showed few control RDG-MPIO bound to the endothelium, indicating that the MPIO-induced hypointensities in these animals do not result from binding to α_v_β_3_. Further histological analysis showed large numbers of iron-laden macrophages and/or microglia, closely associated with blood vessels, leading to the suggestion that the MPIO may be taken up by perivascular macrophages. Whilst the majority of iron detected was visible inside macrophages, single endothelium-bound MPIO were also present in mice injected with RGD-MPIO, with significantly more endothelium-bound MPIO present compared to those given control RDG-MPIO. In accord with the above findings in non-enhancing tumours, these data support the notion that specific binding of RGD-MPIO to angiogenic blood vessels is present.

Anecdotally, unconjugated MPIO have also been found to be retained in tumour bearing striatum, suggesting that the non-specific uptake of MPIO is specific to tumour. However, caution is required in the interpretation of these results, since only a single mouse was tested and the surface of the unconjugated MPIO has different properties and charge to the peptide- and antibody-conjugated MPIO. Non-specific uptake of MPIO has not been previously observed in our experience using antibody targeted MPIO. Therefore, antibody-conjugated MPIO targeting endothelial VCAM-1 were also tested in the 4T1-GFP model to determine whether MPIO with different targeting moieties behave differently *in vivo.* MPIO-induced hypointense voxels were observed in the tumour bearing striatum in mice imaged with VCAM-MPIO, but not in the contralateral striatum, nor in mice imaged with control IgG-MPIO. Further, histological analysis showed marked endothelial VCAM-1 expression at the margins of the tumours, and single MPIO bound to VCAM-1 positive blood vessels. These data are in accord with previous studies in the micrometastatic stages of the 4T1-GFP model^47,49^ and other macrometastasis models^52^ showing that antibody-conjugated MPIO are capable of sensitive and specific imaging of VCAM-1 positive blood vessels. However, neither of the two animals imaged with IgG-MPIO showed any MRI-detectable hypointensities or macrophage uptake by immunohistochemistry, in line with our previous studies. Nor was macrophage uptake of MPIO evident in any of the VCAM-MPIO injected animals. Thus, across a total cohort of 5 mice injected with antibody-conjugated MPIO, no non-specific hypointensities were observed and no macrophage uptake was evident, despite the presence of large, contrast-enhancing tumours in 4/5 animals. These data support the conclusion that there is no non-specific retention of antibody-targeted MPIO. These findings are in contrast to the peptide-targeted MPIO findings discussed above, where animals with large, contrast-enhancing tumours showed significant non-specific retention of both RGD-MPIO and control RDG-MPIO.

Together the above findings suggest that peptide-targeted MPIO, but not antibody-targeted MPIO, are non-specifically retained within the tumour microenvironment, and that this is particularly evident when the BBB is compromised. Given the presence of iron-laden macrophages observed in tumour tissue, we first assessed whether the numbers of these increased in animals injected with peptide-targeted MPIO compared to either those given antibody-targeted MPIO or animals with no exposure to MPIO. We found, however, that late-stage tumours (day 28-35 timepoints) had similar levels of Prussian blue-stained macrophages, regardless of the targeting moiety, and similar levels were also present in tumours with no MPIO exposure. These findings indicated that there is no increase in the absolute number of iron-laden macrophages in the peptide-MPIO injected animals specifically.

As the MPIO-induced hypointensities were only observed in tumours with frank BBB breakdown, we next considered whether peptide-targeted MPIO had simply extravasated into the parenchyma. It is generally thought that MPIO with a diameter of 1 μm are unable to cross the BBB as they are too large to extravasate from the circulation, and are used in preference to iron oxide nanoparticles for this reason. To directly test this assumption, we used a model of CINC-1 induced BBB breakdown. In accord with the above premise, however, only occasional single MPIO and iron-laden macrophages were observed in the CINC-1 injected hemisphere, indicating that neither peptide- nor antibody-targeted MPIO cross a permeable BBB *per se* to any significant degree.

A previous clinical study from Iv *et al.* showed a correlation between macrophage density in high grade glioma with R2* measurements following ferumoxytol administration^60^, suggesting that the non-specific retention of peptide-targeted MPIO in the 4T1-GFP model may be related to the tumour microenvironment, rather than BBB breakdown *per se,* and that factors within the tumour microenvironment may prime macrophages/microglia to take up MPIO across a compromised BBB. Studies by Leftin and Koutcher, demonstrated that iron-laden macrophages in breast cancer brain metastases are pro-inflammatory polarised specifically^58^. Since pro-inflammatory macrophages exhibit increased phagocytic activity, we hypothesise that pro-inflammatory macrophages within the tumour microenvironment, that are already iron-laden and have access to the blood through close association with the BBB, may be predisposed to take up peptide-targeted MPIO, and to a greater degree when the BBB is compromised. Such uptake would lead to an increase in the volume of observed hypointensities, and could explain the non-specific hypointensities seen when using control RDG-MPIO. Currently, endogenous iron and MPIO cannot be distinguished using Prussian blue histology. Future studies may be able to use fluorescently or otherwise labelled beads to distinguish between endogenous iron and MPIO.

Non-specific MPIO induced hypointensities have not been observed in any previous studies using MPIO to study brain pathology^33,47,49,51,61,62^. However, all previous studies have used an antibody as the targeting moiety compared to the peptides used in this study. It is unclear why peptide- rather than antibody-targeted MPIO may be more readily phagocytosed by perivascular macrophages, leading to non-specific contrast effects, but our current data suggest that antibody targeting may be preferable to peptide targeting under these circumstances. At the time of study design, no antibodies targeted to mouse integrin α_v_β_3_ and suitable for *in vivo* use were commercially available. Therefore, a direct comparison of peptide and antibody targeting moieties was not possible in this study, but warrants further study in the future.

## Conclusions

Whilst specific binding of the α_v_β_3_-targeted molecular imaging agent RGD-MPIO was observed, a number of confounds, including pre-existing hypointensities and non-specific retention of peptide-targeted MPIO, need to be overcome before this approach can progress. Little evidence of iron retention in the brain parenchyma, either in macrophages or as bound MPIO, was found in areas of frank BBB breakdown induced by CINC-1 in the absence of a tumour. We propose, therefore, that the non-specific retention of peptide-targeted MPIO is related to uptake by primed perivascular macrophages/microglia within the tumour environment, rather than BBB breakdown *per se.* Whilst low levels of endogenous iron-laden macrophages may reduce the sensitivity of the molecular MRI results, pre-contrast imaging would enable differentiation between endogenous macrophages and specific contrast. Moreover, antibody conjugated MPIO enable sensitive and specific imaging of endothelial targets, with no apparent non-specific retention, and thus show significant advantages over peptide-targeted MPIO. These findings have important implications for the emerging field of molecular MRI, suggesting that antibodies rather than peptides may be preferable as the targeting ligand.

## Acknowledgments

The authors would like to thank the Biomedical Services Unit and the Imaging Core for their assistance with animal and MRI experiments. This work was supported by Cancer Research UK [grant number C5255/A15935]; and the CRUK/EPSRC Cancer Imaging Centre in Oxford [grant number C5255/A16466]. JB is supported by Charlie Perkins, Chevening, and Green Templeton College Scholarships.

## Data Availability Statement

The data that support the findings of this study are available from the corresponding author upon reasonable request.

## Abbreviations

BBB: blood-brain barrier
BSA: bovine serum albumin
CINC-1: cytokine induced neutrophil chemoattractant
fSEMS: fast spin echo multi-slice
MGE3D: multi gradient echo three dimensional
MPIO: microparticles of iron oxide
PBS: phosphate buffered saline
RDG: Arg-Asp-Gly peptide, scrambled control
RGD: Arg-Gly-Asp peptide, targeting integrin α_v_β_3_
SEMS: spin echo multi-slice
USPIO: ultra-small superparamagnetic iron oxide
VCAM-1: vascular cellular adhesion molecule 1

## Notes

### Competing Interest Statement

The authors have declared no competing interest.

