## Supplementary Material for "Imaging angiogenesis in an intracerebrally induced model of brain macrometastasis using α_v_β_3_-targeted iron oxide microparticles"

**Supplementary Information**

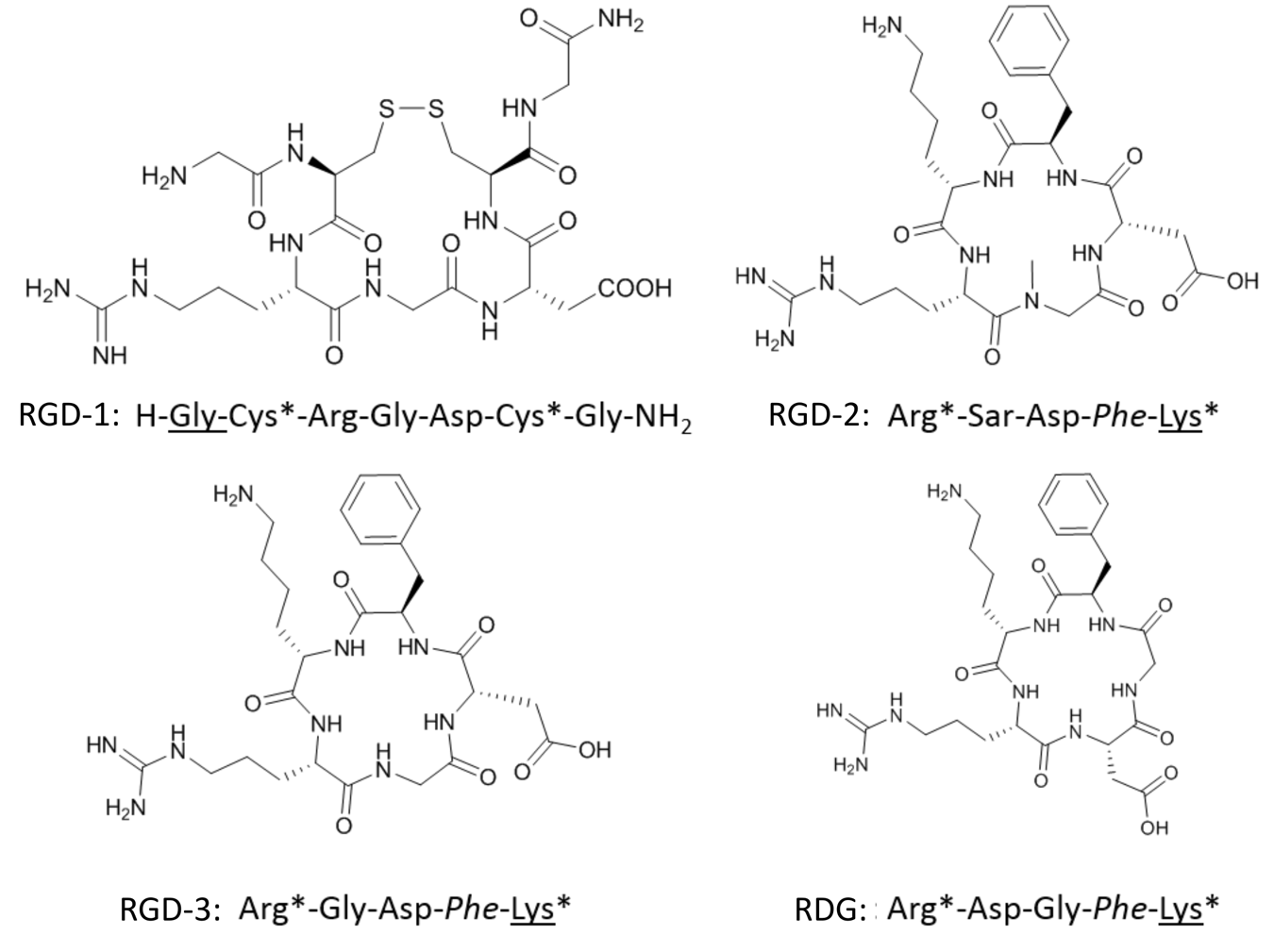

**Supplementary Figure 1:** Chemical structures of the cyclic RGD peptides used in this study. The amino acid underlined is the free amine that binds to the MPIO, those with asterisks indicate the cyclic part of the structure, and those in italics indicate D-amino acids. RGD-1, RGD-2 and RGD-3 are cyclic-RGD structures, while RDG is a scrambled peptide sequence which is used as a negative control. Gly – glycine, Cys – cysteine, Arg – arginine, Asp – aspartic acid, Sar – sarcosine (N-methyl-glycine), Phe – phenylalanine, Lys – lysine.

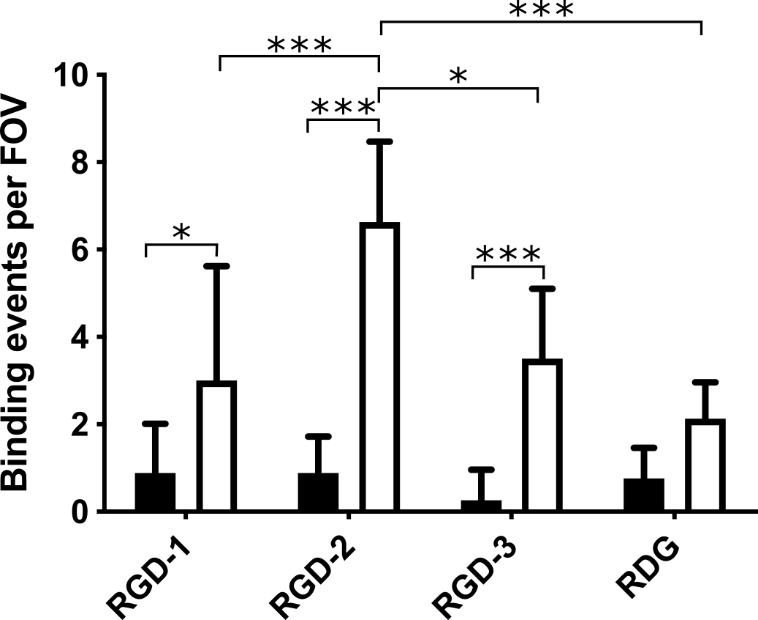

**Supplementary Figure 2:** Binding of cyclic-RGD conjugated MPIO to α_v_β_3_ coated capillaries (white bars) under physiological flow conditions. Binding of MPIO to a BSA coated capillary (black bars) as a negative control was also measured. Error bars represent standard deviation (n = 8 FOVs per capillary and peptide), one-way ANOVA with Sidak’s post-hoc test *p < 0.05, ***p < 0.001.

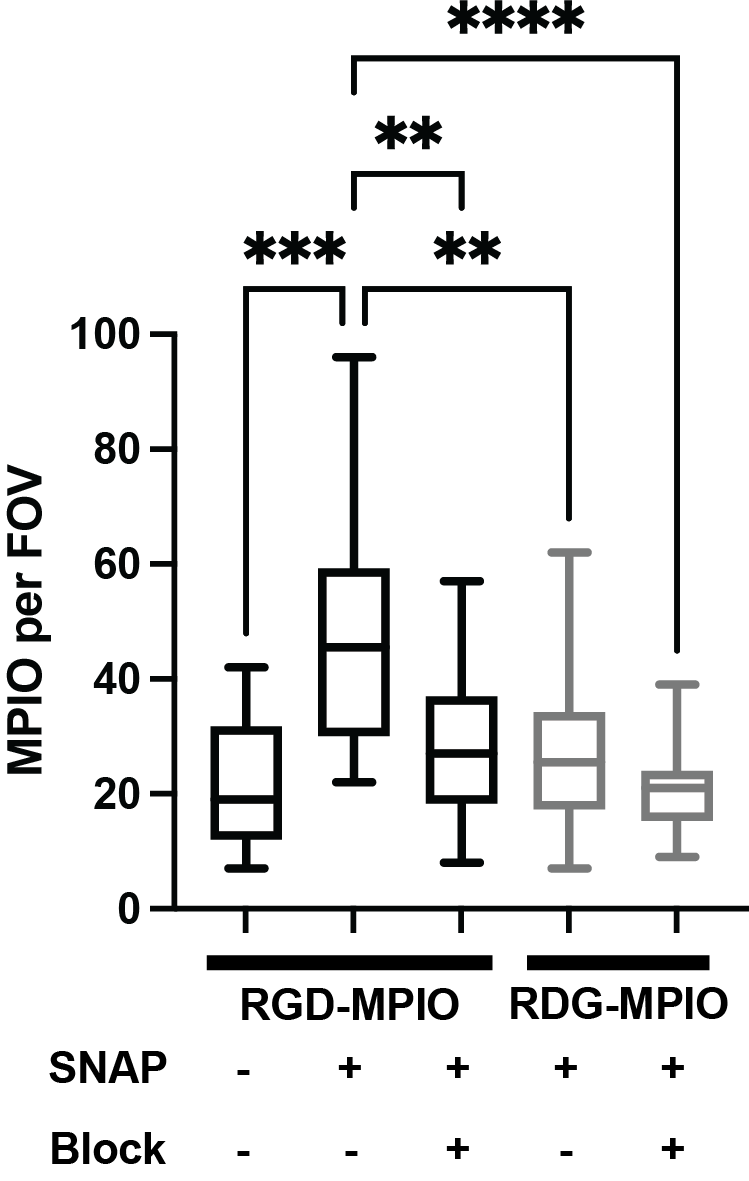

***Supplementary Figure 3:*** RGD-MPIO bind specifically to α_v_β_3_ expressing cells. HUVEC-C cells were treated with S-nitroso-n-acetylpenicillamine (SNAP) to induce α_v_β_3_ expression, followed by a pre-treatment block with either RGD peptide (for cells treated with RGD-MPIO) or control RDG peptide (for cells treated with RDG-MPIO). RGD-MPIO bind specifically to α_v_β_3_ expressing, unblocked cells. Box and whisker plot shows median and interquartile range, Welch’s ANOVA with Dunnett’s multiple comparison test, **p < 0.01, ***p < 0.001, ****p < 0.0001.

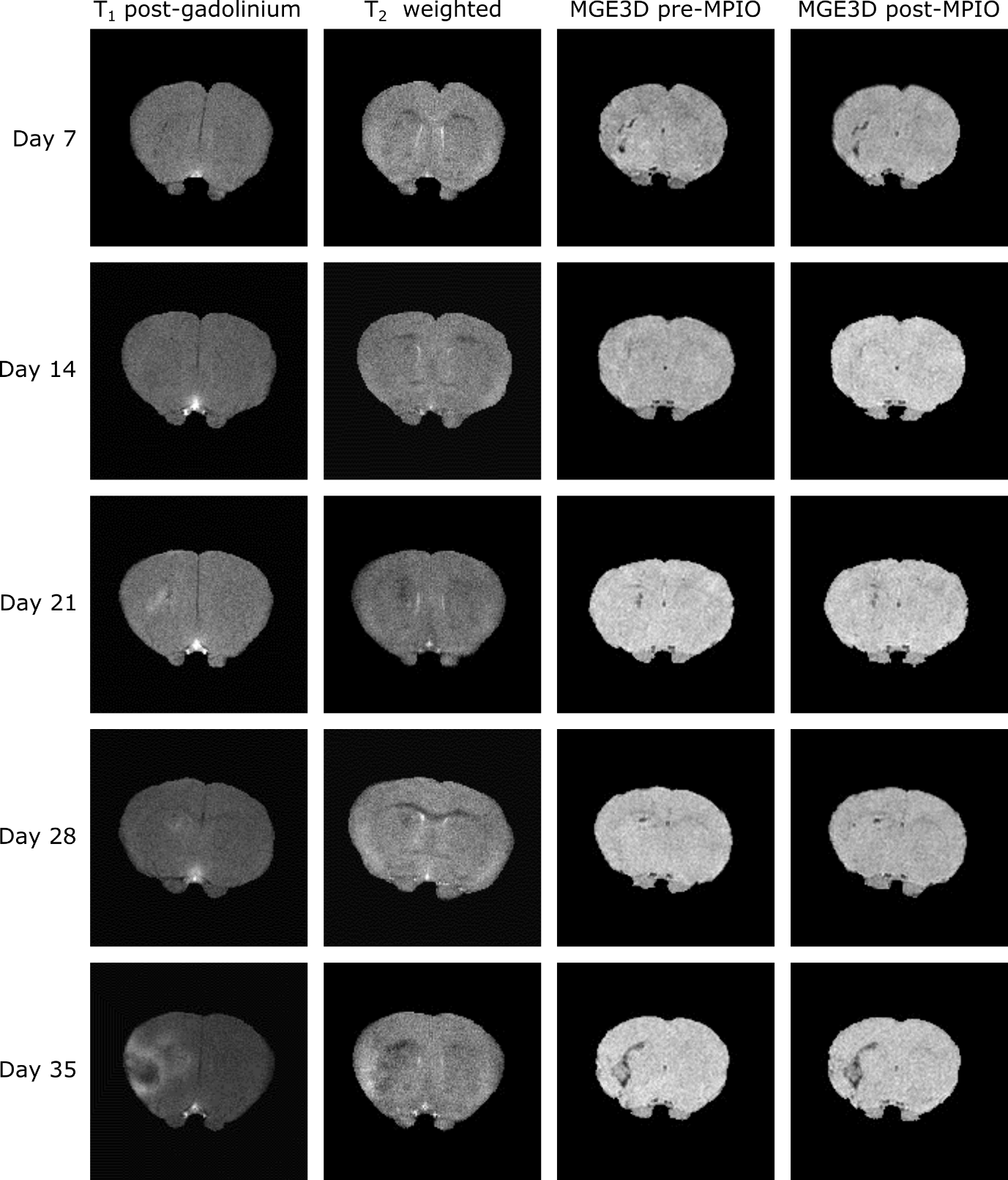

***Supplementary Figure 4:*** Representative images for each time-point in the 4T1-GFP model from mice injected with RGD-MPIO. Each column, from the left, shows T_1_-weighted post-gadolinium images, T_2_-weighted images, pre-RGD-MPIO MGE3D images and post-RGD-MPIO contrast MGE3D images.

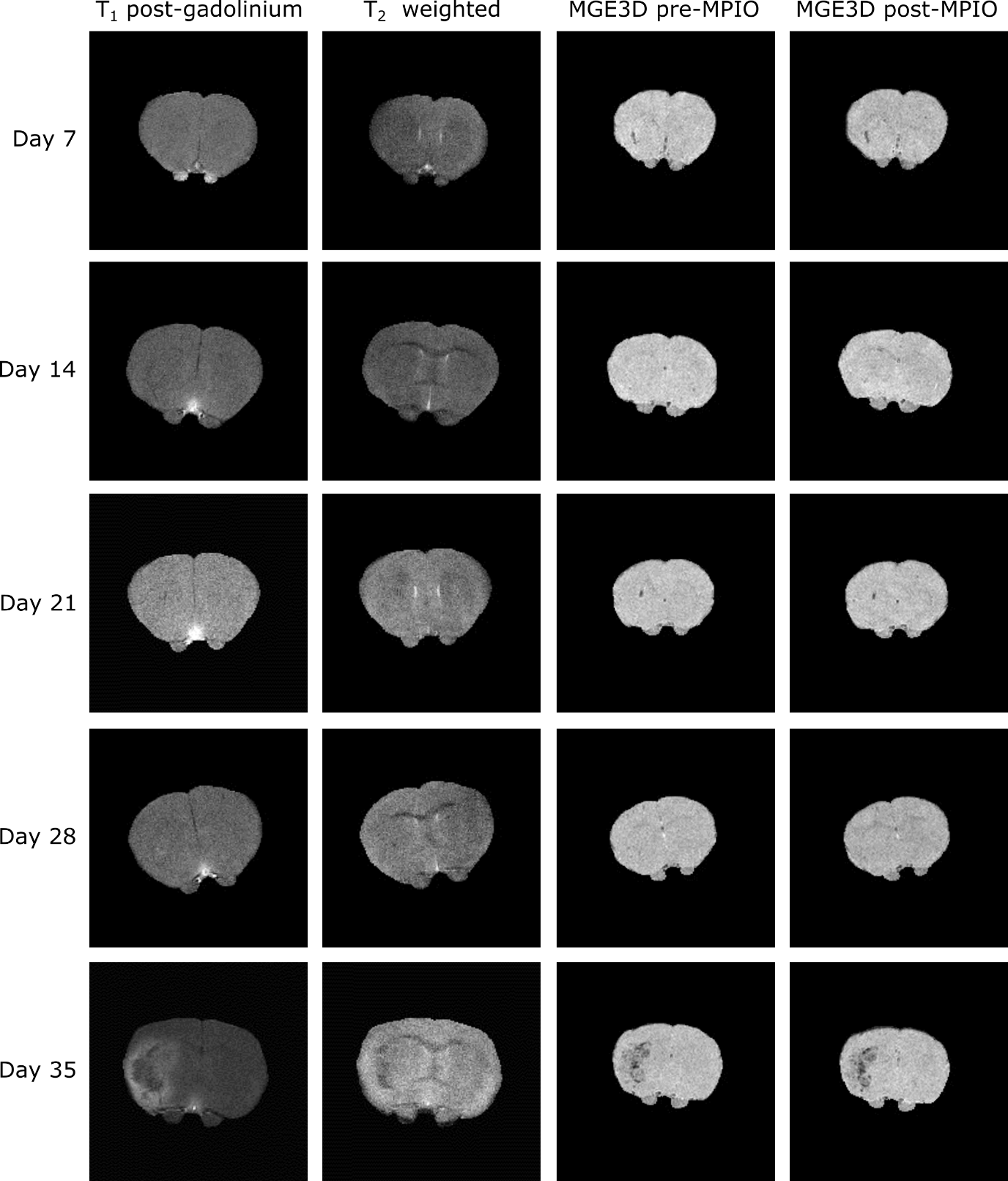

***Supplementary Figure 5:*** Representative images for each timepoint in the 4T1-GFP model from mice injected with control RDG-MPIO. Each column, from the left, shows T_1_-weighted post-gadolinium images, T_2_-weighted images, pre-RGD-MPIO MGE3D images and post-RGD-MPIO contrast MGE3D images.

***Supplementary Table 1:*** Numbers of gadolinium enhancing and non-enhancing tumours at each timepoint.

| **MPIO type** | **Timepoint** | **Non-enhancing** | **Enhancing** | **Total** |
| --- | --- | --- | --- | --- |
| **RGD-MPIO** | Day 7 | 3 | 0 | 3 |
|  | Day 14 | 2 | 0 | 2 |
|  | Day 21 | 1 | 2 | 3 |
|  | Day 28 | 1 | 3 | 4 |
|  | Day 35 | 0 | 5 | 5 |
|  | **Totals** | **7** | **10** | **17** |
| **RDG-MPIO** | Day 7 | 3 | 0 | 3 |
|  | Day 14 | 3 | 0 | 3 |
|  | Day 21 | 3 | 0 | 3 |
|  | Day 28 | 3 | 1 | 4 |
|  | Day 35 | 1 | 2 | 3 |
|  | **Totals** | **13** | **3** | **16** |

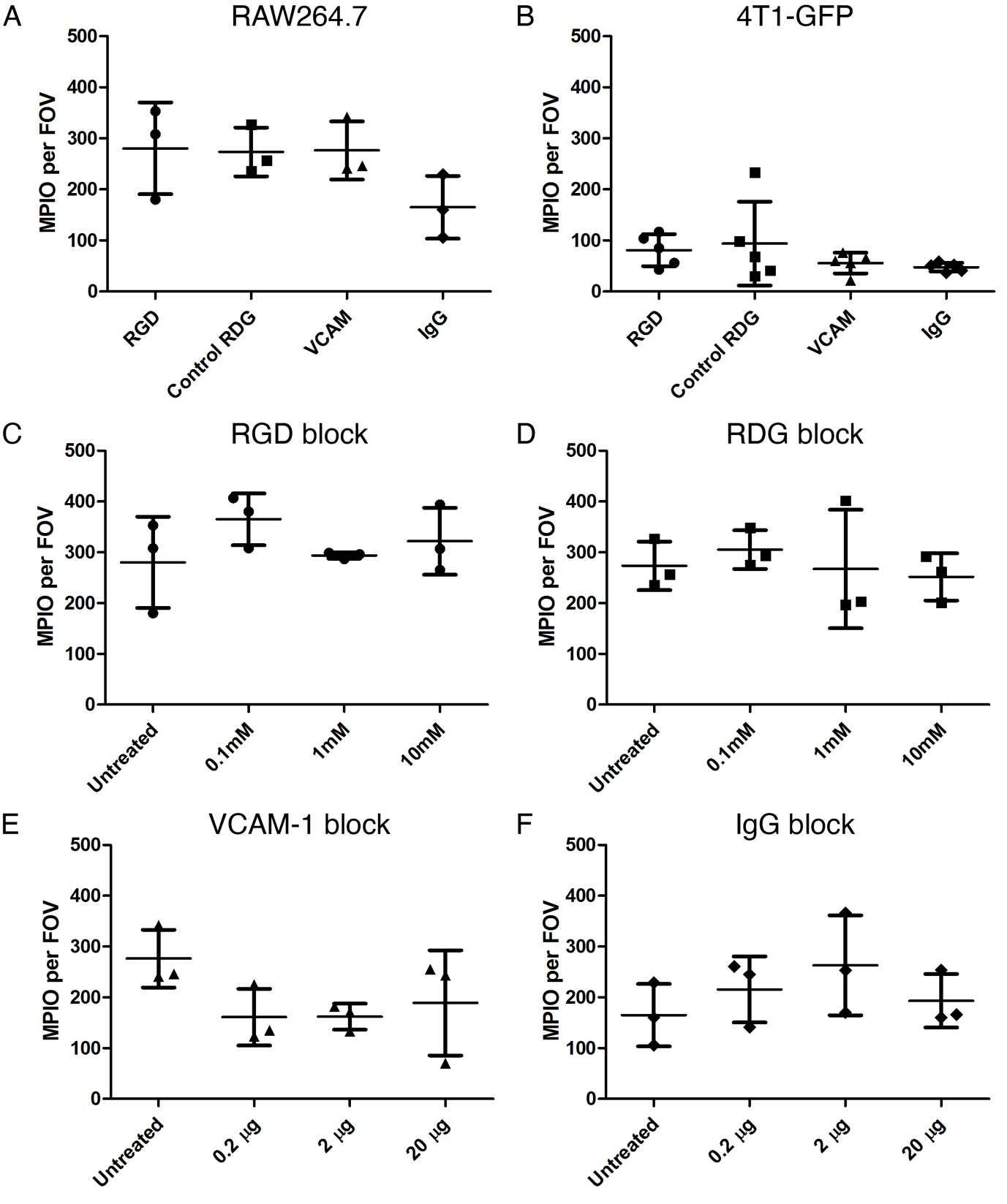

***Supplementary Figure 6:*** Uptake of MPIO *in vitro* by 4T1-GFP tumor cells and macrophages. (A) Uptake of all four types of MPIO (RGD, control RDG, VCAM, IgG) by RAW 264.7 macrophages. (B) Uptake of all four types of MPIO by 4T1-GFP tumour cells. Uptake was significantly higher in macrophages than for 4T1-GFP cells (2-way ANOVA, p < 0.001) for all MPIO types (Bonferroni post-hoc tests p < 0.05). (C) Uptake of RGD-MPIO by RAW264.7 macrophages following pre-treatment with increasing concentrations of RGD peptide. (D) Uptake of control RDG-MPIO by RAW264.7 macrophages following pre-treatment with increasing concentrations of control RDG peptide. (E) Uptake of VCAM-MPIO by RAW264.7 macrophages following pre-treatment with increasing concentrations of VCAM-1 antibody. (F) Uptake of IgG-MPIO by RAW264.7 macrophages following pre-treatment with increasing concentrations of IgG antibody. Data are shown as individual data points (n = 3 per group), with error bars representing mean ± standard deviation.

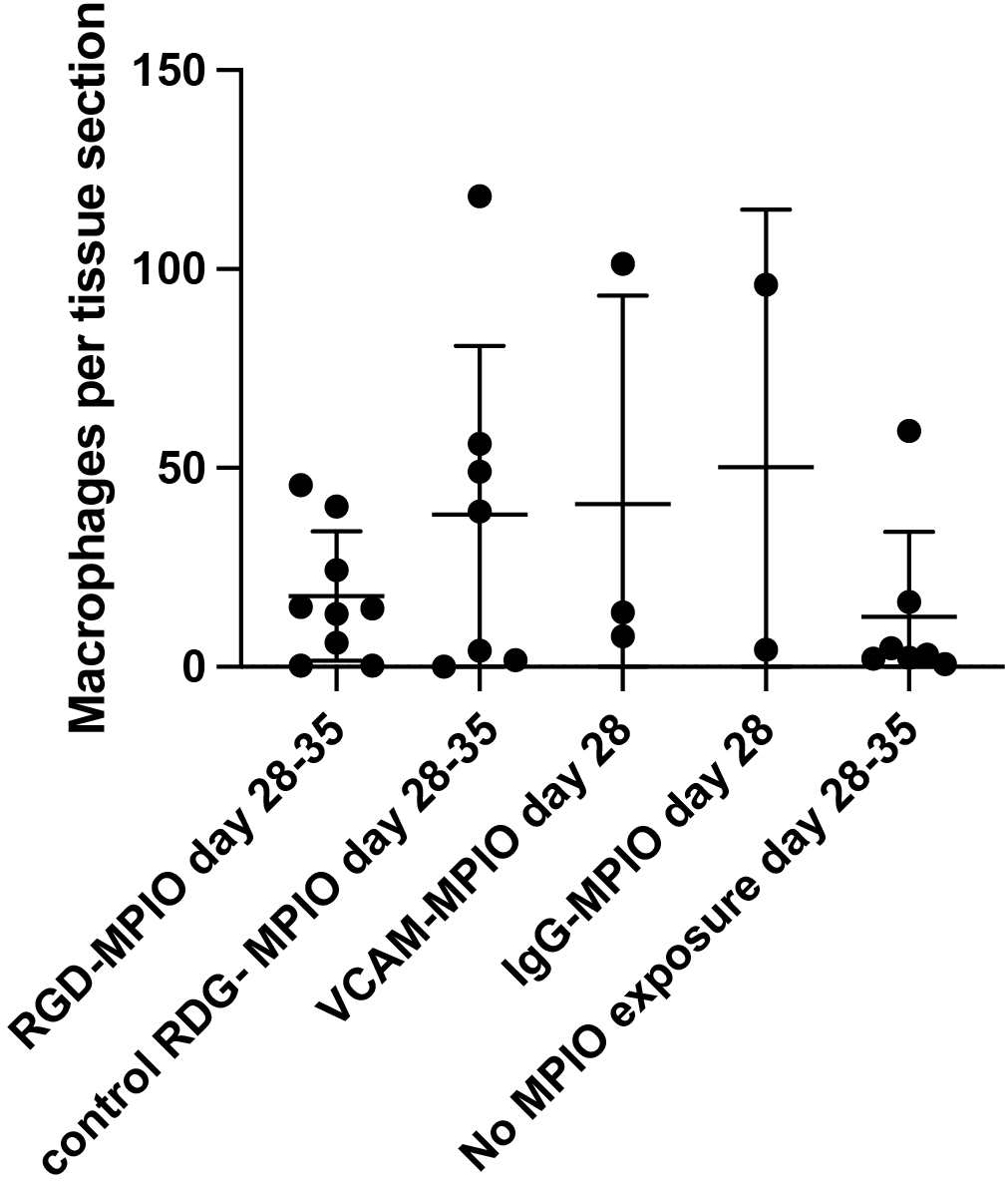

***Supplementary Figure 7:*** Iron-laden macrophages present in Prussian blue stained histology sections in the 4T1-GFP tumour model. No significant differences were present between groups. Error bars represent standard deviation.

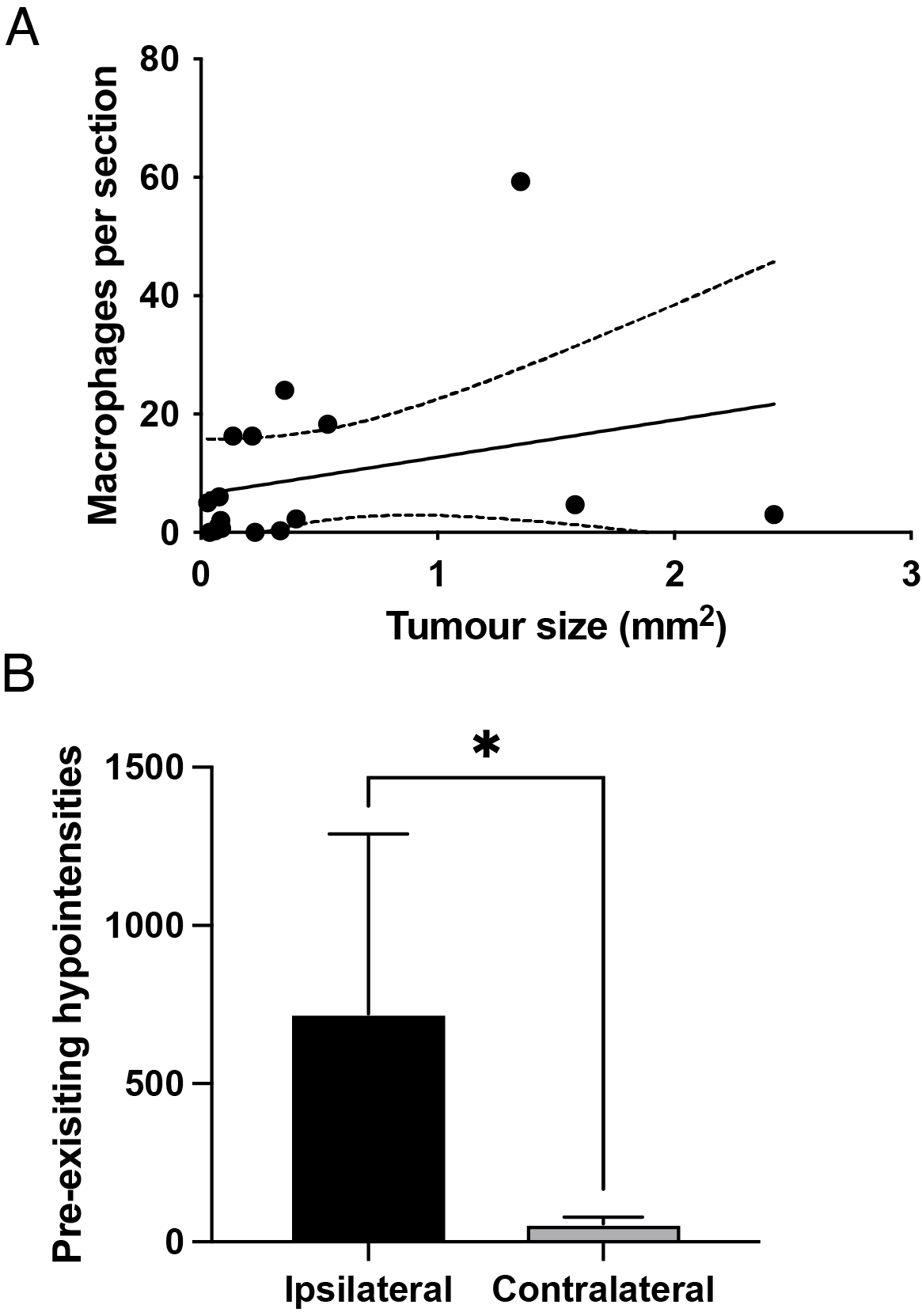

***Supplementary Figure 8:*** Endogenous iron laden macrophages cause pre-existing hypointensities in mice with no exposure to MPIO. (A) The number of endogenous iron-laden macrophages per tissue section does not correlate with tumour size (p > 0.2, Pearson’s coefficient R^2^ = 0.08). (B) Pre-existing hypointense voxels are observed in the tumor-bearing hemisphere. Graph showing pre-existing hypointense voxels in 4T1-GFP tumors at the day 28 timepoint, in mice with no exposure to MPIO. Significantly more hypointense voxels are present in the tumor bearing ipsilateral striatum than the contralateral striatum (p < 0.05, paired t-test, n = 6). Error bars represent mean ± standard deviation. *p < 0.05

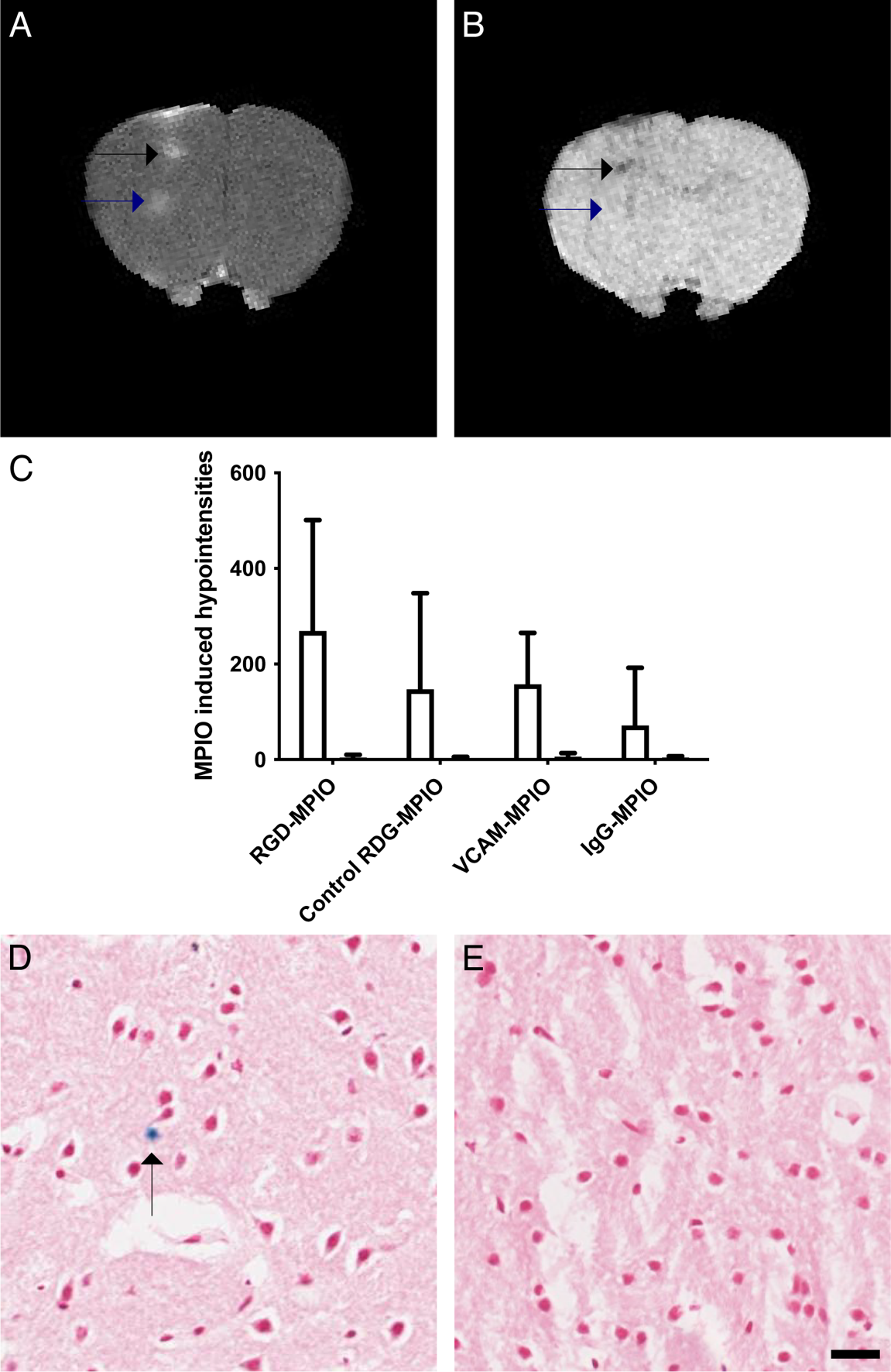

***Supplementary Figure 9:*** MPIO uptake in the brain following CINC-1 induced BBB breakdown. (A) Gadolinium enhanced T_1_-weighted scan showing frank BBB breakdown (indicated by arrows). (B) MPIO induced hypointense voxels in a post RGD-MPIO injection MGE3D scan are visible in some, but not all areas in which BBB breakdown has occurred (indicated by arrows). (C) Quantitation of MPIO induced hypointense voxels, for the ipsilateral striatum (white bars) and the contralateral striatum (black bars). Bars represent mean ± standard deviation. No significant differences were found between treatment groups. (D) Histological analysis showed no evidence of tissue damage, and very few iron aggregations were present in the injected striatum (indicated by arrow). (E) No evidence of iron aggregations was found in the contralateral striatum. Scale bar = 20 µm.
